# Restriction of the growth and biofilm formation of *ESKAPE* pathogens by caprine gut-derived probiotic bacteria

**DOI:** 10.1101/2023.06.30.546197

**Authors:** Prerna Saini, Repally Ayyanna, Rishi Kumar, Sayan Kumar Bhowmick, Bappaditya Dey

**Author notes:** **Corresponding address:** Bappaditya Dey, Scientist-E, National Institute of Animal Biotechnology (NIAB), Survey No. 37, Extended Q City Road, Gowlidoddi, Gachibowli, Hyderabad, Telangana, India 500032. Gut probiotics limit ESKAPE pathogen growth and biofilms.

## Abstract

The accelerated rise of antimicrobial resistance (AMR) poses a significant global health risk, necessitating the exploration of alternative strategies for combating pathogenic infections. Biofilm-related infections, which are unresponsive to standard antibiotics, often require the use of higher-order antimicrobials with toxic side effects and a potential for disrupting the microbiome. Probiotic therapy, with its diverse benefits and inherent safety, is emerging as a promising approach for preventing and treating various infections and as an alternative to antibiotic therapy. In this study, we isolated novel probiotic bacteria from the gut of domestic goats (*Capra hircus*) and evaluated their antimicrobial and antibiofilm activities against the ‘ESKAPE’ group of pathogens. We performed comprehensive microbiological, biochemical, and molecular characterizations, including analysis of the 16S-rRNA gene V1-V3 region and the 16S-23S ISR region, on 20 caprine gut-derived lactic acid bacteria (LAB). Among these, six selected LABs demonstrated substantial biofilm formation in anaerobic conditions, and exhibited robust cell surface hydrophobicity and autoaggregation properties highlighting their superior enteric colonization capability. Notably, these LAB isolates exhibited broad-spectrum growth inhibitory and anti-biofilm properties against ‘ESKAPE’ pathogens. Additionally, the LAB isolates were susceptible to antibiotics listed by the European Food Safety Authority (EFSA), within the prescribed Minimum Inhibitory Concentration limits, suggesting their safety as feed additives. The remarkable probiotic characteristics exhibited by the caprine gut-derived LAB isolates in this study strongly endorse their potential as compelling alternatives to antibiotics and as direct-fed microbial (DFM) feed supplements in the livestock industry, addressing the escalating need for antibiotic-free animal products.

## INTRODUCTION

Antimicrobial resistance (AMR) is a growing public health concern globally, as many pathogens are becoming resistant to standard antibiotics. The ‘*ESKAPE*’ group of six nosocomial pathogens is leading the priority pathogen list of multidrug-resistant (MDR) and extensively drug-resistant (XDR) bacteria that includes, i.e., *Enterococcus faecium, Staphylococcus aureus, Klebsiella pneumoniae, Acinetobacter baumannii, Pseudomonas aeruginosa, and Enterobacter spp.* [1]. These pathogens are capable of ‘escaping’ the bactericidal action of various antimicrobial agents. Inappropriate use or overuse of antibiotics has resulted in a global emergence and spread of these pathogens, causing outbreaks, community-acquired infections, and transmission by the foodborne and waterborne routes to animals and humans. Infection-related fatalities caused by drug-resistant (DR) pathogens are expected to account for the highest number of deaths worldwide by 2050 [2]. Biofilm formation is a major mechanism that DR and MDR-ESKAPE bacteria use to exhibit drug resistance phenotype [3]. Biofilms protect specialized dormant persister cells that are tolerant to antibiotics as well as host immune cells, leading to difficult-to-treat recalcitrant infections [4]. Antibiotics are given alone or in combination to effectively treat such infections. However, with every passing year, the number of antibiotics to treat these infections is declining, predisposing humanity toward a future with fewer antibiotics that will probably become ineffective in the near future [5]. Hence, there is a dire need to find safe and natural antibiotic alternative agents, such as probiotics, to treat infections caused by such pathogens [3, 6].

Probiotics are live microorganisms that confer health benefits on the host when administered in adequate amounts and are considered a potential alternative to antibiotics (FAO/WHO, 2001) [7]. Most probiotics belong to Lactic acid bacteria (LAB), a group of bacteria with a generally regarded as safe (GRAS) status, and are the oldest known probiotic to inhibit or treat infections caused by DR pathogens [8, 9]. LAB strains belonging to *Lactobacillus and Bifidobacterium* genera are known to inhibit pathogens by a plethora of mechanisms, including a competitive exclusion for binding sites, adhesion to the intestinal mucosa, host immunomodulation, enhancement of epithelial barrier integrity, production of organic acids, hydrogen peroxide, bacteriocins, and antimicrobial peptides, etc. [10]. The application of LAB has shown promise in treating infections by *ESKAPE* bacteria in both animals and humans. For example, topical application of *L. acidophilus or L. reuteri* was effective in treating wound infections in mice caused by *A. baumanii* [11]. The application of *L. fermentum* improved the condition of ischemic wounds in rabbits [12]. Similar activity against skin pathogens such as *E. coli, P. aeruginosa, S. aureus,* and *Propionibacterium* have been reported for *L. plantarum ATCC 10241* and *L. delbrueckii DSMZ 20081* [13, 14]. In addition*, L. casei subsp. rhamnosus, L. fermentum, L. acidophilus, and L. plantarum* all prevented the adhesion and regrowth of *E. faecalis and E. faecium* biofilms [15]. *Lactobacillus gasseri* LBM220 isolated from the feces of breastfed infants showed strong antibacterial activity against all six MDR- *ESKAPE* pathogens [16]. LAB can inhibit the growth of a variety of livestock pathogens, including bovine mastitis-causing *Methicillin Resistant Staphylococcus aureus* (MRSA), which also causes skin abscesses and septicemia [17, 18]. Further, gram-negative bacillus *K. pneumoniae*, a common causative agent of clinical mastitis in dairy cattle, is an emerging zoonotic and foodborne pathogen worldwide [19, 20]. *Lactobacillus plantarum CIRM653* was able to impair *K. pneumoniae*- biofilms independently of its bactericidal effect [21]. *L. delbrueckii subsp. delbrueckii* LDD01 also showed the best inhibitory results against *K. pneumoniae* [22]. *Pseudomonas aeruginosa* is also associated with many diseases in livestock and companion animals, including urinary tract infections in dogs, mastitis in dairy cows, and endometritis in horses [23, 24]. *Lactobacillus fermentum* isolates showed growth inhibition and anti-biofilm effects against all MDR, XDR, and pan-drug-resistant (PDR) *P. aeruginosa* strains [25]. Notably, the *L. acidophilus* ATCC 4356 strain is well-known for its strong biofilm growth inhibition against a majority of the *P. aeruginosa* strains [26, 27]. EFSA has listed *E. faecalis* as an opportunistic pathogen for birds, poultry, and reptiles [28]. *Enterococcus faecalis* has also been detected in animals, meat, and meat-based products, as well as from human fecal samples and patients with bloodstream infections [29]. Bio surfactants and conditioned medium from several probiotic bacteria were found to prevent adhesion and biofilm formation by *E. faecalis* [30]. Lipoteichoic acids from *L. plantarum* were proven effective in disrupting mature *E. faecalis* biofilms [31]. Further, livestock animals are the main reservoir of Shiga toxin-producing *E. coli* with zoonotic potential, and several LAB isolates exhibited antagonistic activity against these *E. coli* strains [32, 33].

Since the antibacterial properties of probiotics LABs depend on strain specificity, comprehensive characterization and evaluation of probiotic properties of new or novel LAB isolates from natural sources are essential before selecting them for preclinical evaluation for their potential use [34, 35]. To date, there is no study demonstrating the isolation and inhibitory effect of caprine gut-derived *LAB* against the *ESKAPE* group pathogens. Here, we isolated LAB from the small intestine of domestic goat, identified by biochemical and molecular methods, assessed the probiotic properties, such as acid and bile tolerance, surface properties, intrinsic colonization-cum-biofilm formation ability, antibiotics susceptibility, and finally demonstrated the growth-antagonistic, and anti-biofilm effect against *ESKAPE* group pathogens.

## RESULTS

### Isolation, identification, and biochemical characterization of LAB from Caprine gut

Gastrointestinal (GI) infections in livestock ruminants not only have significant implications for animal health, and productivity, but they also impact public health as a source of zoonotic enteric and other diseases in humans. Resident probiotic bacteria in the small intestine of domestic goat may have evolved niche-specific colonization and host-favoring properties that promote the maintenance of enteric homeostasis and antagonizes the growth of pathobionts in the gut. To utilize the beneficial properties of such enteric probiotic LAB, first, we collected small intestinal (jejunum) tissues from goats and subjected them to a standard MRS media-based LAB isolation method [36, 37]. Following several rounds of isolation of potential-LAB colonies with creamy white color on MRS agar, we selected 54 colonies for further confirmation by *Lactobacillus* genus-specific PCR using 16S-rRNA gene-specific primers, Forward-R16-1 and Reverse-LbLMA1 as described previously by Dubernet and colleagues (**Supplementary Table-1**) [38]. Out of 54 colonies, we selected 20 colonies that showed an expected amplification of ∼250 bp **(Supplementary Fig. 1a)**. Further, these 20 isolates were subjected to a second round of PCR and amplicon sequencing targeting 16S-rRNA V1-V3 region and 16S-23S Intergenic Spacer Region (ISR) for species identification [39–42]. Two separate primer pair sets were used to amplify ∼509 bp **(Supplementary Fig. 1b)** and ∼565 bp regions **(Supplementary Fig. 1c)** from the 16S-rRNA-V1-V3 region and 16S-23S ISR using 16S(8-27)-F and V3(519-536)-R, and 16S(ISR)-F and 23S(ISR)-R primer pairs, respectively (**Supplementary Table-1**). Out of the 20 LAB-like colonies that were subjected to PCR, followed by Sanger sequencing and NCBI-BLAST-based nucleotide homology analysis, 12 isolates showed the closest identity to *Lactobacillus sp*., while the other colonies were found to belong to either *Enterococcus spp*. or *Acetobacter spp*. (Table-1). All the sequences were submitted to NCBI **Bio-project No. PRJNA985412, PRJNA986841, PRJNA986842**. As for the 13 *Lactobacillus sp*. species, 5 of them were identified as *L. plantarum*, 2 colonies were *L. salivarius*, 2 were *L. crispatus*, 2 were *L. amylovorous*, 1 was *L. kitasatonis*, and another was *L. jhonsonii* (Table-1). Subsequently, phylogenetic analysis of the 12 LAB isolates using either of the sequences of 16S rRNA-V1-V3 region (**Fig.1a**) or 16S-23S ISR (**Fig.1b**) via hierarchical clustering using the Neighbour Joining method exhibited close evolutionary relationship and relatedness amongst the caprine gut-derived LAB isolates. As shown in the dendrograms in **Fig.1**, isolates belonging to the same species of *Lactobacillus* were mapped as closely related by both the target sequences. Moreover, isolates from the same animal origin exhibited the closest phylogenetic relationship.

**Fig. 1:**
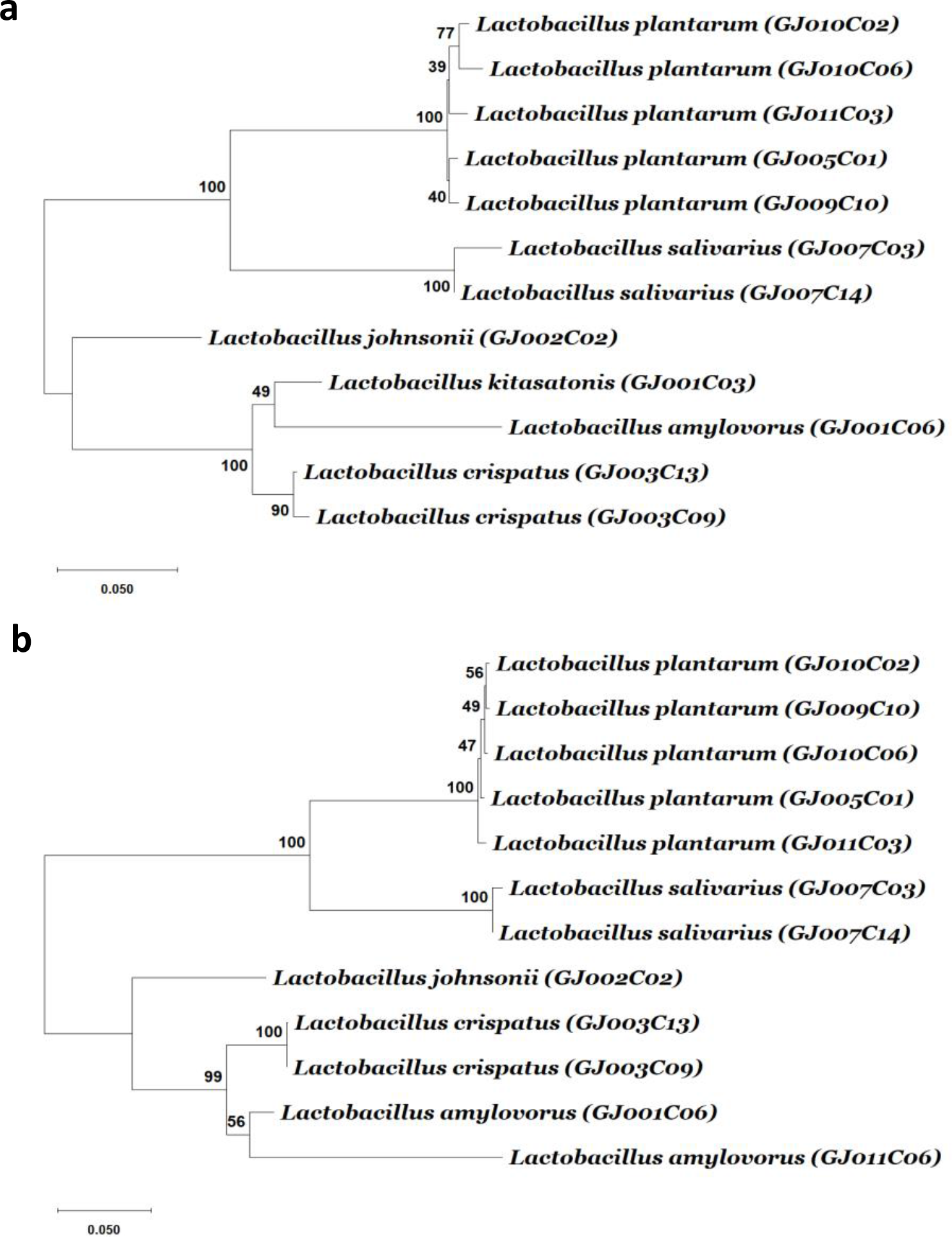
The dendrogram tree of LAB isolates based on 16S rRNA gene V1-V3 region and 16S-23S ISR sequences. The amplicons from the 16S rRNA gene V1-V3 region and 16S-23S ISR were sequenced using the primers 16S(8-27)-F and 16S(ISR)-F, respectively. From each isolate, either of the sequences was aligned using ClustalW with default parameters. The alignment results were then used to find out best fitted nucleotide-substitutional model. The Kimura 2-parameter model and Tamura 3-parameter model are used for the V1-V3 region, and ISR region, respectively. The neighbor-joining method (bootstrap test -1000 replicates) was used to generate the hierarchical clustering-based dendrogram trees based on (**a**) 16S rRNA gene V1-V3 region, and (b) 16S-23S ISR region. A unit of the distance scale represents the percentage of differences between two sequences. Evolutionary analyses were conducted using MEGA11 software.

**Table- 1:**
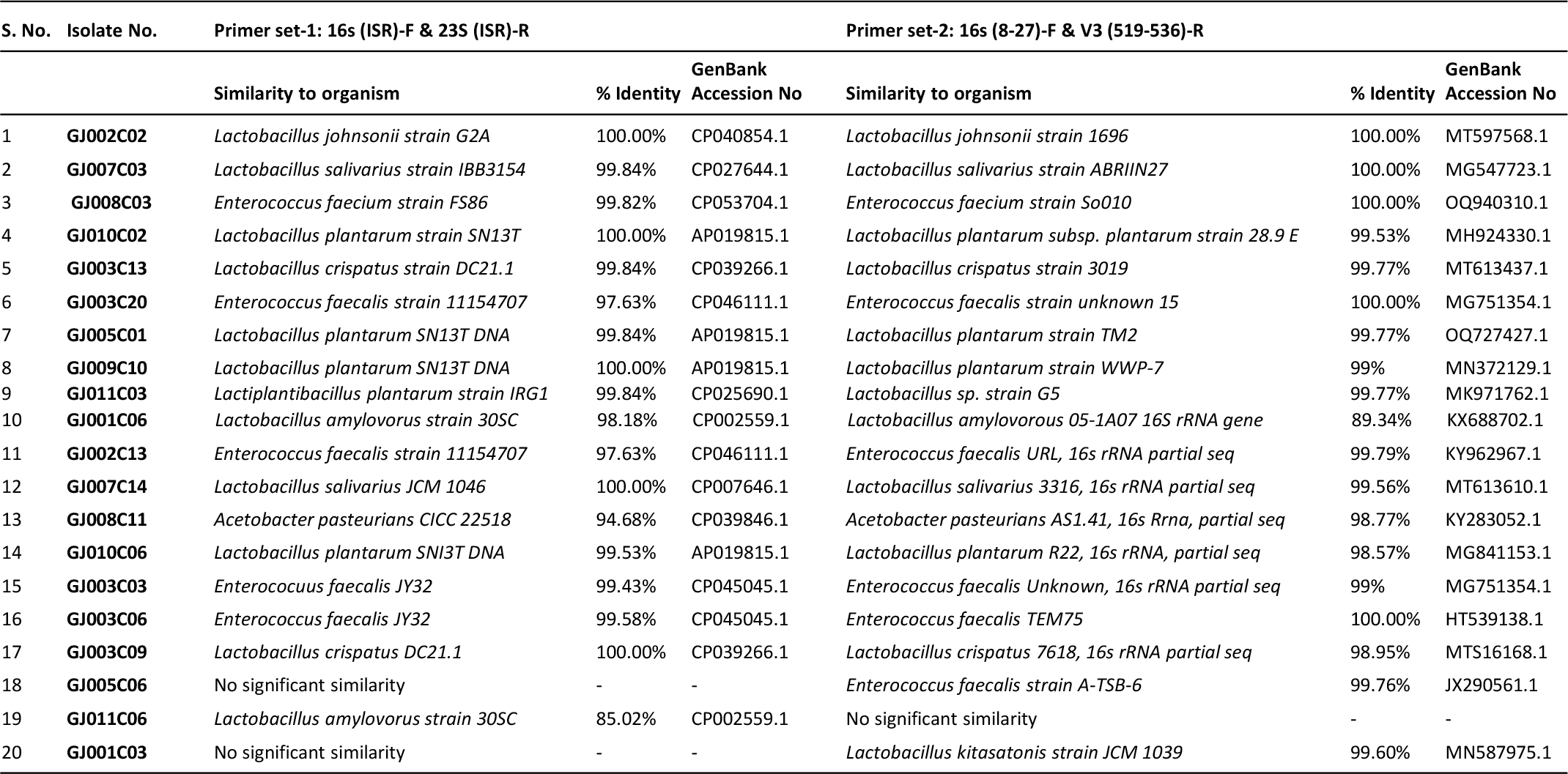
16s-rRNA gene V1-V3 region and 16s-23s ISR amplicon sequencing and identification of LAB isolates. For LAB species identification PCR was performed targeting the 16S-rRNA V1-V3 region and 16S-23S Intergenic Spacer Region (ISR), and two separate primer pair amplified ∼509 bp, and ∼565 bp. The amplified PCR products were purified and sequenced via Sanger sequencing using the forward primers of each PCR described above. The trimmed sequences were subjected to the NCBI-BLAST analysis with default parameters for identifying the closest bacterial species with maximum percentage identity.

Further, 12 selected LAB isolates were subjected to a series of biochemical tests to meet the criteria for LAB species as described in *Bergey’s Manual of Determinative Bacteriology* [43]. These include Gram staining, catalase test, nitrate reduction (NR) test, indole test, and Vogues Proskauer (VP) test. All 12 isolates showed Gram-positive staining, of which 6 were identified as catalase negative while the remaining 6 were catalase positive. All 12 LAB isolates showed negative test results for the indole test, nitrate test, and VP test (except one showing weak + VP test) (Table-2). These features of the LAB isolates corroborate well with the standard characteristics of *Lactobacillus spp*. As Catalase-negative probiotics are considered superior as they can survive and thrive in the low-oxygen environment of the digestive system, and are also less likely to cause harmful changes in the gut microbiota by producing reactive oxygen species (ROS), which can damage the intestinal lining and trigger inflammation, 6 catalase negative LAB isolates were selected for further characterization by carbohydrate fermentation test.

**Table- 2.** Biochemical characterization of LAB isolates. Each LAB isolate was screened for Gram staining, catalase activity, indole test, for the production of indole from tryptophan; nitrate test for the reduction of nitrate to nitrite; and Vogues Proskauer (VP) test to determine the presence of acetyl methyl carbinol after glucose fermentation. Six of the Gram-positive and catalase-negative LAB isolates were selected for further analysis.

| S. No. | LAB isolates | Gram's stain | Catalase test | Nitrate test | Indole test | VP test |
| --- | --- | --- | --- | --- | --- | --- |
| 1. | *L. kitasatonis* (GJ001C03) | + | + | - | - | - |
| 2. | *L. amylovorus* (GJ001C06) | + | + | - | - | weak + |
| 3. | *L. jhonsonii* (GJ002C02) | + | + | - | - | - |
| 4. | *L. crispatus* (GJ003C09) | + | - | - | - | - |
| 5. | *L. crispatus* (GJ003C13) | + | + | - | - | - |
| 6. | *L. plantarum* (GJ005C01) | + | - | - | - | - |
| 7. | *L. salivarius* (GJ007C03) | + | - | - | - | - |
| 8. | *L. salivarius* (GJ007C14) | + | + | - | - | - |
| 9. | *L. plantarum* (GJ009C10) | + | - | - | - | - |
| 10. | *L. plantarum* (GJ010C06) | + | - | - | - | - |
| 11. | *L. plantarum* (GJ011C03) | + | - | - | - | - |
| 12. | *L. amylovorus* (GJ011C06) | + | + | - | - | - |

+, positive for the given test; -, Negative for the given test; weak +, slight positive reaction.

Probiotic lactobacillus with complex sugar fermentation capability is important for their use in ruminant feed because ruminants rely on microbial fermentation of complex sugars in their digestive process [44]. Here, we assessed the carbohydrate fermentation ability of these 6 LAB isolates on 10 different carbohydrate sugars, i.e., monosaccharides (Xylose, Arabinose, Galactose, Mannose), disaccharides (Cellobiose, Maltose, Melibiose, Sucrose, Trehalose), and trisaccharide (Raffinose) using a Hilacto^TM^ Identification kit for *Lactobacillus* that also includes catalase and esculin test. All 6 LAB isolates utilized sugars at different rates of hydrolysis, indicating their varied carbohydrate fermentation capability (Table-3**, Supplementary Fig.2**). Further, the LAB isolates displayed a positive esculin test, which is a typical feature of *Lactobacillus spp.* confirming their ability to break down complex coumarin glycoside-esculin into glucose and esculetin due to the presence of β-glucosidase or esculinase enzyme [45].

**Table- 3.**
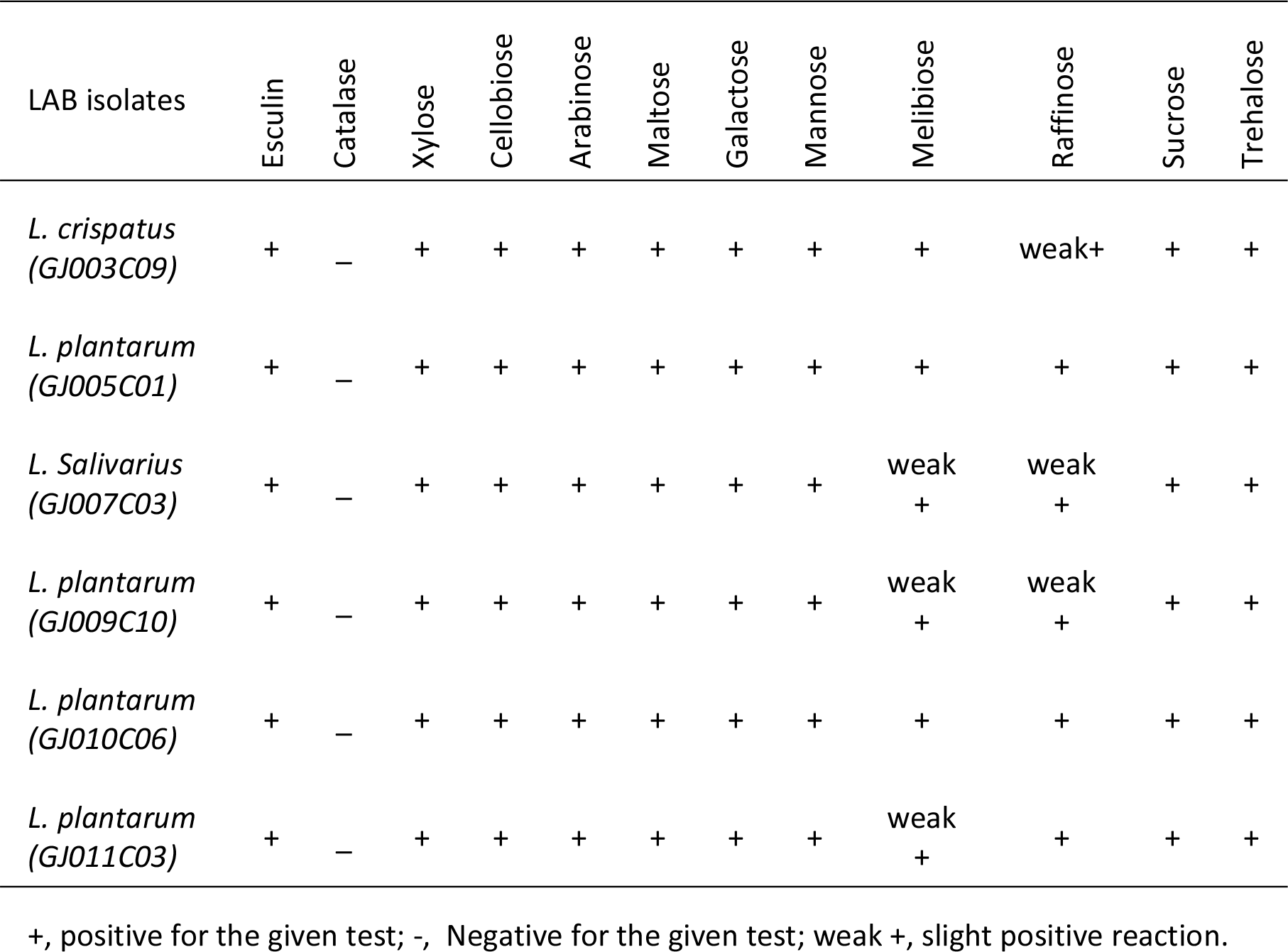
LAB isolates exhibit differential carbohydrate fermentation ability. The table depicts the carbohydrate fermentation ability of selected 6 LAB isolates. The carbohydrate fermentation was measured via colorimetric identification based on the principle of pH change and substrate utilization for the genus *Lactobacillus*. All isolates were found esculin positive, catalase-negative, and capable of fermenting different carbohydrate sugars with varied abilities.

### Caprine gut-derived LAB isolates exhibited acid stress and bile salt tolerance

The ability to tolerate harsh conditions, like low pH or acidic environment in the stomach, and sustenance in the high bile salts concentration at the initial part of the small intestine are the major determinants for the survival of probiotic bacteria in the gastrointestinal tract [46]. Each of the 6 selected LAB isolates displayed different degrees of sensitivity towards pH and bile salt stress, as evidenced by their differential growth pattern monitored over a period of 12 hours at different pH (2.0, 3.0, 4.0, and 6.5) and bile salt (0.5%, 1.0%, and 2.0%) containing media. In a non-acidic environment at pH 6.5, 4 of the LAB isolates exhibited typical sigmoidal bacterial growth, except GJ010C06 and GJ003C09 isolates that displayed longer lag phases with minimal growth (**Fig. 2a**). At pH 4.0, half of the LAB isolates exhibited moderate growth, while other halves displayed reduced or stalled growth (**Fig. 2b**). Nevertheless, all the isolates, including GJ010C06 and GJ003C09, survived acid stress at lower pH of 2.0 and 3.0, while GJ011C03 strain exhibited the highest growth at these low pH environments (**Fig. 2c, d**).

**Fig. 2.**
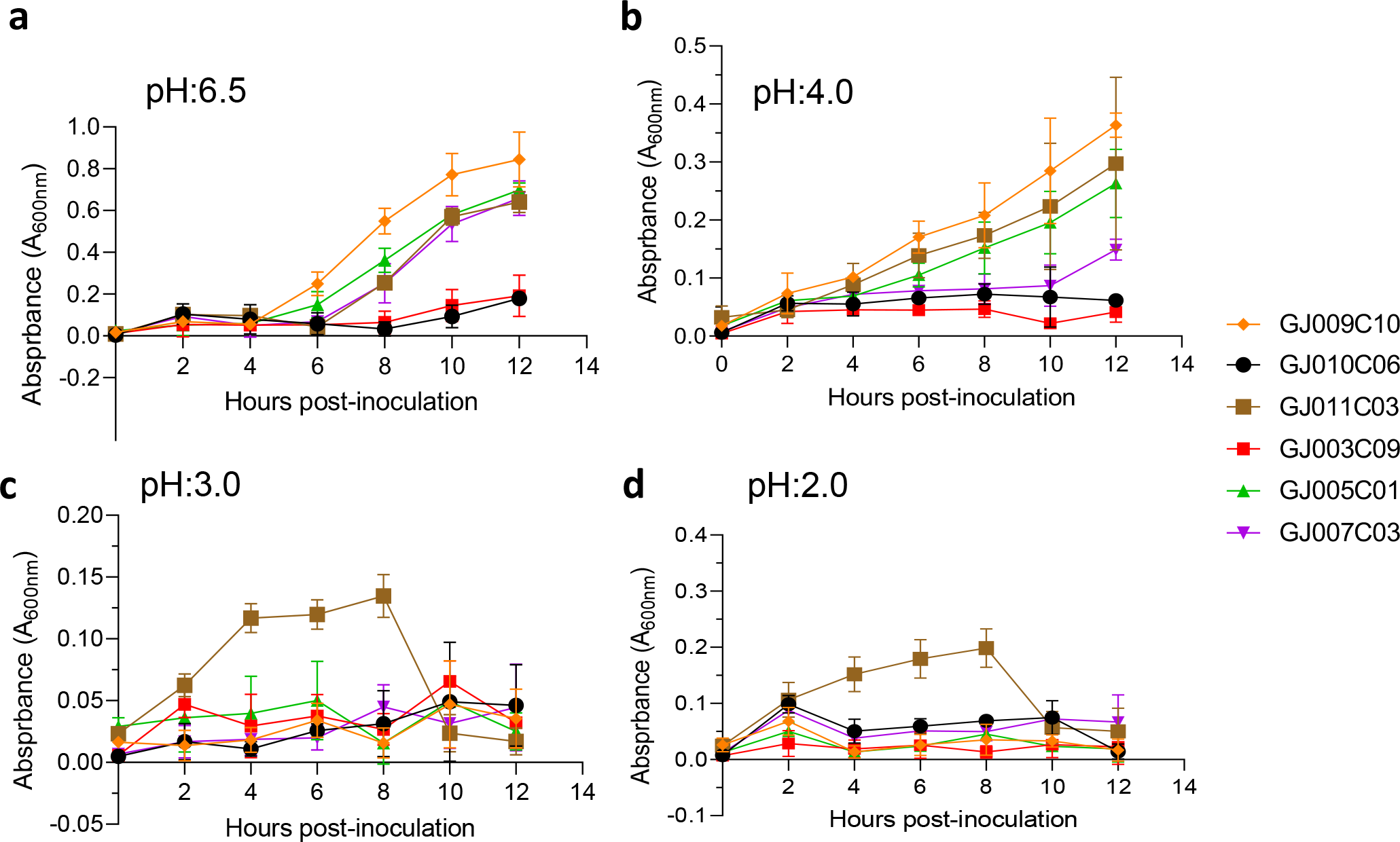
Profiling of the ability of LAB isolates to pH stress. Tolerance of LAB isolates to lower pH was assessed in three different conditions in addition to the MRS media (pH 6.5). Absorbance (A_600nm_) was taken at an interval of 2 hours and till 12 hours post-inoculation. Each connecting point in the line graphs depicts the mean ± SD of the absorbance values at each time point indicating the growth of LAB isolates at (a) pH 6.5, (b) pH 4.0, (c) pH 3.0, and (d) pH 2.0 conditions.

When bile salts were added to the growth medium at three different concentrations (0.5%, 1%, and 2% W/V), all selected LAB isolates exhibited suppressed growth compared to the standard growth media (**Fig. 3a**). However, they were able to endure the salt stress for up to 12 hours in media containing 0.5 % bile salt (**Fig. 3b**). In case of 1.0 % and 2.0 % bile salt containing media, after an initial normal growth till 4 hours post-inoculation, the LAB isolates displayed a sharp reduction in culture absorbance, followed by a plateaued phase of growth (**Fig. 3c & d**). Among the 6 LAB isolates, GJ009C10, GJ010C06, and GJ011C03 maintained higher culture absorbance levels compared to the other three LAB isolates.

**Fig. 3.**
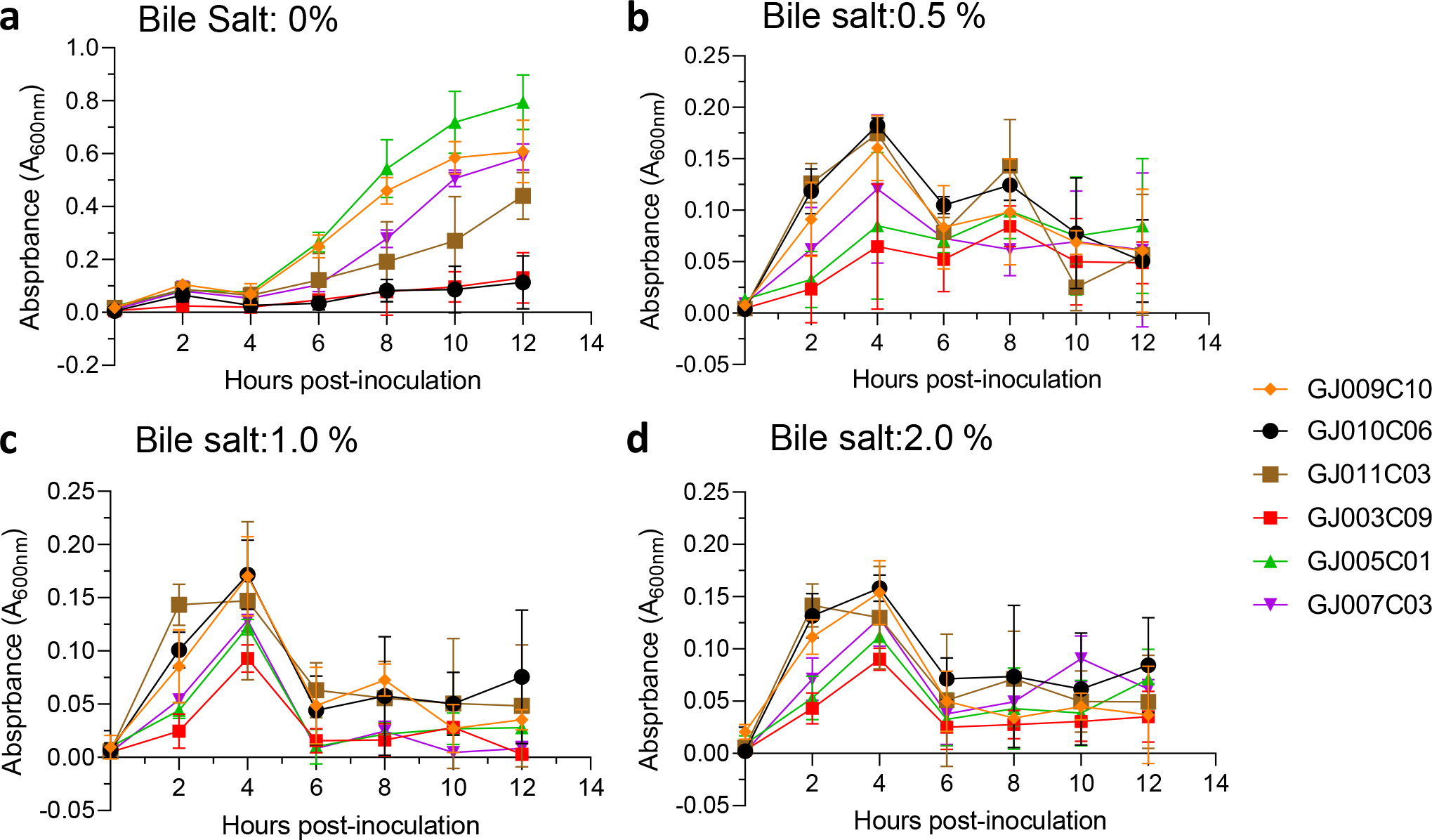
Bile tolerance profile of LAB isolates. Bile salt tolerance of LAB isolates was assessed in three different concentrations in addition to the standard MRS media. Absorbance (A_600nm_) was taken at an interval of 2 hours and till 12 hours post-inoculation. Each connecting point in the line graphs depicts the mean ± SD of the absorbance values at each time point indicating the growth of LAB isolates in (**a**) 0% bile salt, (**b**) 0.5%, (**c**)1%, and (**d**) 2% w/v) conditions.

### Cell surface hydrophobicity and auto-aggregation properties of LAB isolates

Probiotic bacteria need to stick to the mucus in the gut to survive, and their ability to do so is crucial for competition with harmful bacteria [47, 48]. Adhesion to the intestinal wall involves both non-specific and specific interactions facilitated by different cell components [49, 50]. The cell surface hydrophobicity, measured by the microbial adhesion to hydrocarbon method (MATH), is an important factor for adhesion capacity and is considered a pre-test for epithelial cell adhesion ability [51].

Similarly, auto-aggregation is a process in which bacteria physically interact with each other with the help of cell surface components such as proteins, carbohydrates, and lipoteichoic acid. Auto-aggregation of probiotics is necessary for adherence to the gut lining and is a barrier against undesirable bacteria [52]. These characteristics provide advantages for probiotics in colonizing the gut. We assessed hydrophobicity using two organic compounds, Xylene and Hexane. All of the LAB isolates exhibited significant adhesion rates (% Hydrophobicity) of over 60%. LAB isolates GJ009C10 and GJ005C01 exhibited >80% adhesion rates (Table-4). Although both the organic solvent-based techniques showed a comparable trend in % hydrophobicity, the xylene-based method yielded higher values compared to the Hexane-based method. The selected LAB isolates also displayed a recommended range of 60% to 80% auto-aggregation percentage, with GJ007C03 and GJ010C06 as top performers (Table-4).

**Table- 4.** Cell surface properties of selected LAB isolates. The table depicts the % Hydrophobicity and % Autoaggregation of LAB isolates. The % hydrophobicity was measured using the microbial adhesion to hydrocarbons (MATH) using hexane, and xylene as organic solvents as described in the Materials & Methods section. Autoaggregation is measured by evaluating the physical clumping ability of the LAB isolates in PBS solution as described earlier. The data is the Mean ± SD of triplicate experiments.

| S. No. | LAB isolates | % Hydrophobicity |  | % Auto-aggregation |
| --- | --- | --- | --- | --- |
|  |  | Xylene | Hexane |  |
| 1. | *L. crispatus* (GJ003C09) | 68.20 ± 0.34 | 68.99 ± 0.79 | 71.52 ± 0.27 |
| 2. | *L. plantarum* (GJ005C01) | 83.63 ± 1.68 | 76.93 ± 1.23 | 70.15 ± 0.77 |
| 3. | *L. salivarius* (GJ007C03) | 69.52 ± 0.51 | 68.68 ± 0.40 | 76.31 ± 0.25 |
| 4. | *L. plantarum* (GJ009C10) | 88.81 ± 2.15 | 84.33 ± 0.63 | 70.82 ± 0.10 |
| 5. | *L. plantarum* (GJ010C06) | 72.73 ± 1.11 | 73.26 ± 0.17 | 75.11 ± 0.15 |
| 6. | *L. plantarum* (GJ011C03) | 83.44 ± 0.77 | 80.46 ± 3.45 | 70.90 ± 0.53 |

### LAB isolates formed more biofilms in the anaerobic environment

A very promising approach for the control of pathogenic bacterial biofilm is the use of probiotics to colonize epithelial surfaces and counteract the proliferation of other bacterial species via competitive exclusion [53]. Probiotic biofilms can promote their self-colonization, longer persistence on the intestinal mucosa, and *Lactobacillus* species produce more robust biofilms than other species [54]. Here, we tested the biofilm formation ability of LAB isolates in both aerobic **(Fig. 4a)** and anaerobic conditions **(Fig. 4b)** as they are facultative anaerobic in nature, and both conditions prevail in the gastrointestinal tract. While all selected LAB isolates formed biofilms under both aerobic and anaerobic conditions, a significantly greater amount of biofilms were formed under anaerobic conditions (**Fig. 4**). The top biofilm-forming LAB isolates under both aerobic and anaerobic conditions were GJ005C01, GJ009C10, GJ003C09, and GJ011C03.

**Fig. 4.**
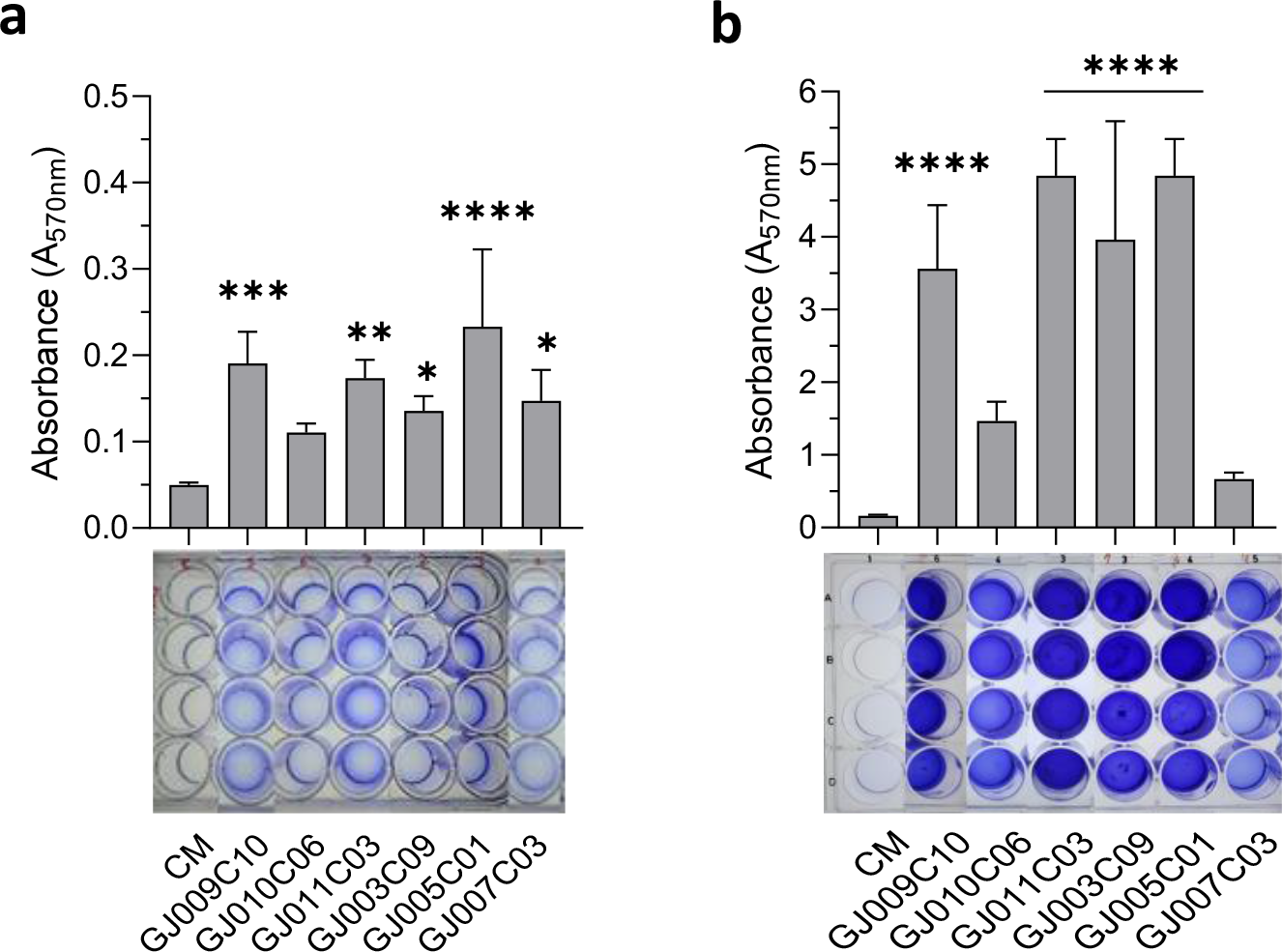
Biofilm formation by LAB isolates in aerobic and anaerobic environments. The biofilm formation ability of LAB isolates was assessed using both **(a)** aerobic, and **(b)** anaerobic growth conditions. The assay was performed in 24-well polystyrene plates. LAB isolates were cultured in static conditions in MRS media at 37 °C for 72 hours. Subsequently, quantitative measurement of biofilm was performed using the standard crystal violet-based assay. The bar diagrams depict the Mean ± SD of the absorbance (A_570nm_) values from triplicate wells. The experiment was performed twice. One-way ANOVA was performed to compare the mean differences of each LAB isolate compared to the base value (control media, CM). *, p<.05, **, p<.01, ***, p<.001, ****, p<.0001. All LAB isolates produced more biofilm in the anaerobic conditions compared to the aerobic environment.

### Caprine gut-derived LAB isolates are non-hemolytic and susceptible to a panel of major antibiotics

While LAB belongs to the GRAS category of bacteria and is typically safe for consumption, newly isolated LAB strains must be proven to be devoid of any toxic effects, such as hemolytic properties, which can lead to serious health problems, including anemia and kidney damage [55, 56]. To ensure that the caprine gut-derived novel LAB isolates are safe, we performed hemolytic profiling for α-hemolysis **(Supplementary Fig. 3a)** and β-hemolysis **(Supplementary Fig. 3b)**, wherein *Streptococcus pneumoniae* and *Staphylococcus aureus* were taken as positive controls for α- and β- hemolysis, respectively [57]. None of the selected caprine gut-derived novel LAB isolates showed a hemolytic effect highlighting their safety for consumption **(Supplementary Fig. 3c).**

Antimicrobial resistance is a significant safety concern when evaluating the use of LAB as a feed additive and therapeutic [58]. The food chain has been identified as a major route of drug-resistant bacterial transmission between animals and humans, highlighting the need to monitor the safety of LAB used in animal nutrition [59]. Here, we screened the caprine gut-derived LAB isolates against nine antibiotics including ampicillin (0.125-64 mg/L), erythromycin (0.0039-2 mg/L), clindamycin (0.0156-8 mg/L), vancomycin (0.125-64 mg/L), gentamicin (0.031-16 mg/L), Kanamycin (0.125-64 mg/L), streptomycin (0.125-64 mg/L), tetracycline (0.125-64 mg/L), and chloramphenicol (0.125-64mg/L) by broth dilution-based MIC method in LSM media using the microbiological breakpoint (BP), or cut-off values recommended by FEEDAP document, and EFSA [60]. The MIC for all the LAB isolates was measured using a U-bottom microplate-based growth assay followed by visual observation at 24 hours post inoculation and were listed in Table-5. All isolated LAB strains were found to be susceptible to nine of the antibiotics tested except vancomycin, and MIC was below the epidemiological cut-off suggested by EFSA [60]. While vancomycin did not have suggested breakpoints by EFSA for *L. plantarum* obligate-, facultative-, heterofermentative and homofermentative species, most of the LAB isolates showed intrinsic resistance to it. Although, screening for streptomycin MIC wasn’t essential for homofermentative *Lactobacillus spp*. [60], as an added caution, we evaluated the MIC for streptomycin and were found to be below the cut-off values provided by EFSA Table-5.

**Table- 5.**
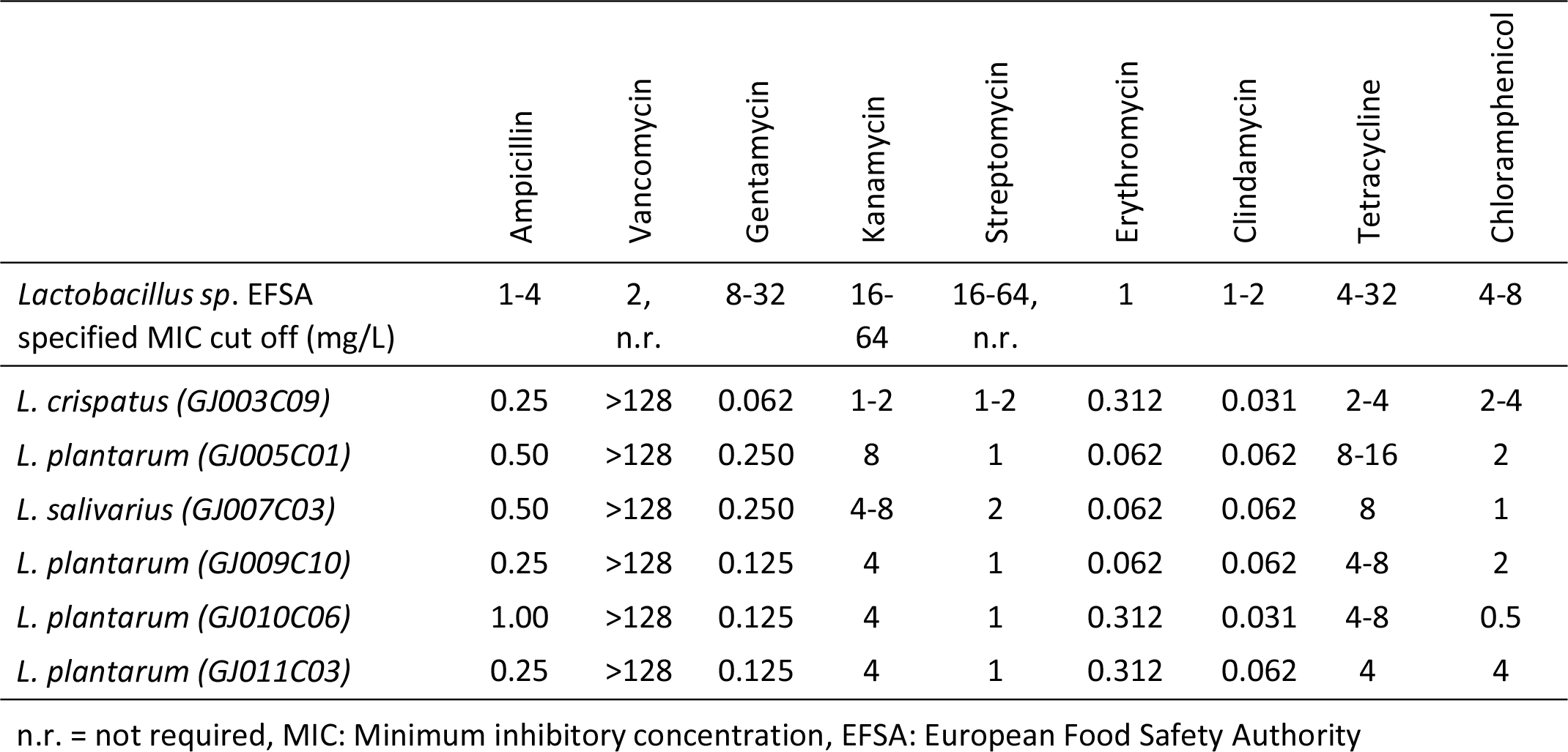
Susceptibility of LAB isolates to EFSA-specified list of antibiotics. The antibiotic susceptibility of the *LAB i*solates was tested for 9 antibiotics using broth micro-dilution method as recommended by FEEDAP and EFSA panel for human and veterinary importance. MIC values of each antibiotic were visually evaluated at 24 and 48 hours as the lowest antibiotic concentrations at which no growth was observed. Experiments were performed thrice.

### Probiotic LAB isolates limit proliferation and biofilm formation by *ESKAPE* pathogens

The *ESKAPE* pathogens, including *E. faecium, S. aureus, K. pneumoniae, A. baumannii, P. aeruginosa, and E. coli* are major causes of infection in animals and humans, but incidences of antibiotics failure against these pathogens is a critical public health problem [61]. These bacteria form biofilms, making them resistant to antibiotics and immune cells. Probiotic LAB is known to inhibit the growth of pathobionts via the production of antimicrobial substances like organic acids, hydrogen peroxide, bacteriocins, antimicrobial peptides, etc. [62]. Antagonistic activity against such pathogens is a prerequisite of a potential probiotic. Therefore, we screened the anti-*ESKAPE* pathogen activity of LAB isolates via two sets of experiments. First, the antagonistic growth potential of LAB- cell-free supernatant (CFS) and cell lysate was evaluated by agar well diffusion assay, and second, anti-biofilm efficacy via measuring their ability to prevent biofilm formation in a microplate-based assay. In this study, six LAB isolates were assessed for antimicrobial activity against six *ESKAPE* pathogens, with results presented in **Fig. 5** and Table-6. Of the isolates, GJ009C10, GJ011C03, GJ005C01, and GJ007C03 demonstrated considerable levels of inhibition against a majority of the *ESKAPE* pathogens. By contrast, growth inhibition was not evident for isolates GJ003C09 and GJ010C06. Results showed similar trends between the CFS and cell-lysate treatments (**Fig. 5**).

**Fig. 5.**
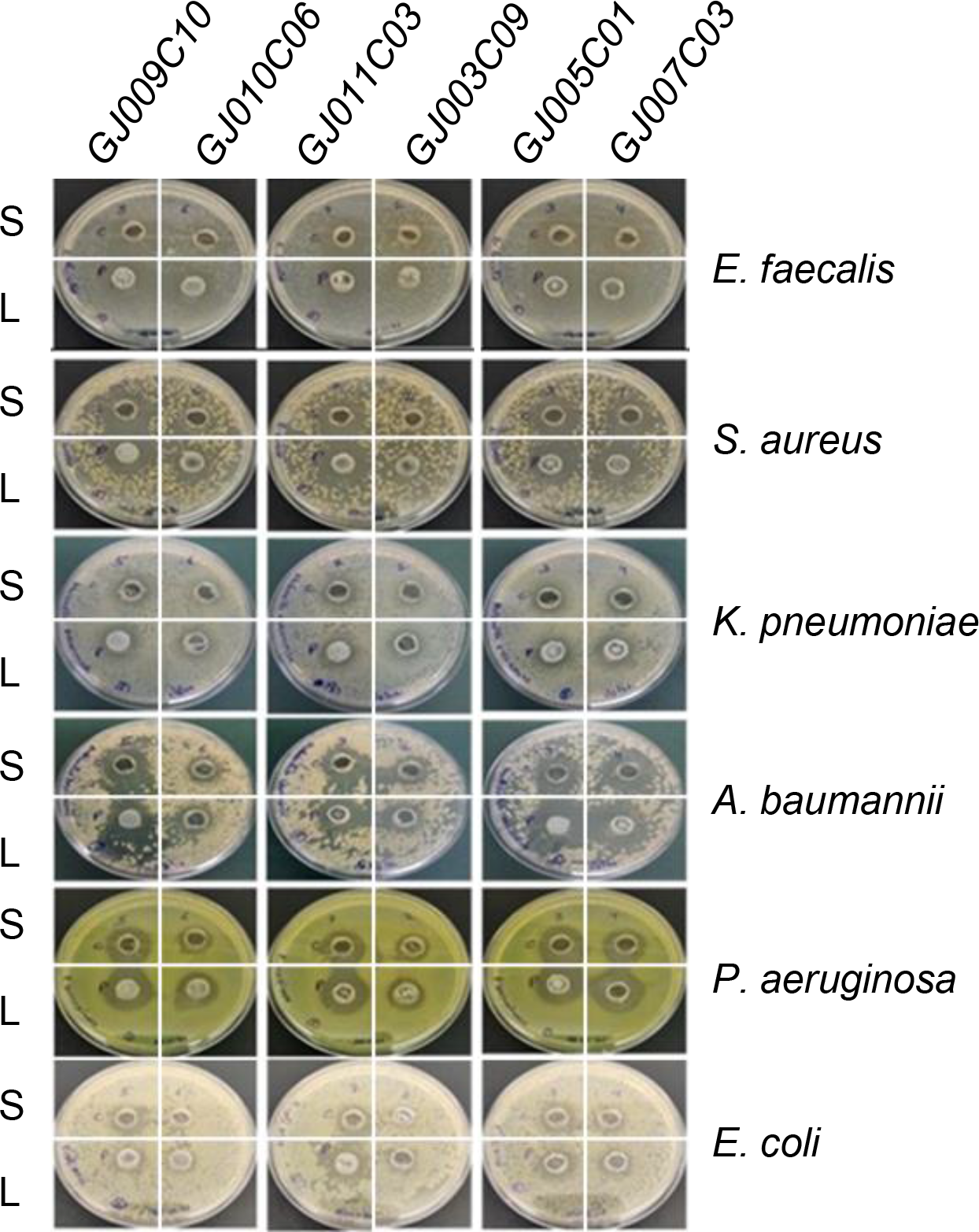
LAB isolates restrict the growth of *ESKAPE* pathogens. The figure depicts representative images of nutrient agar plates from agar well diffusion assay-based antimicrobial activity assay. Each *ESKAPE* bacterial inoculum was spread onto nutrient agar plates. Subsequently, 10 mm diameter wells were made on the agar plates using 1 ml sterile pipette tip bottoms. Sterile-filtered cell-free culture supernatants (S) and cell lysate (L) of LAB isolates were added to each well and incubated for 24 hours at 37°C in aerobic conditions. The zone of inhibition was recorded with the zone of inhibition scale. Experiments are performed thrice.

**Table- 6.**
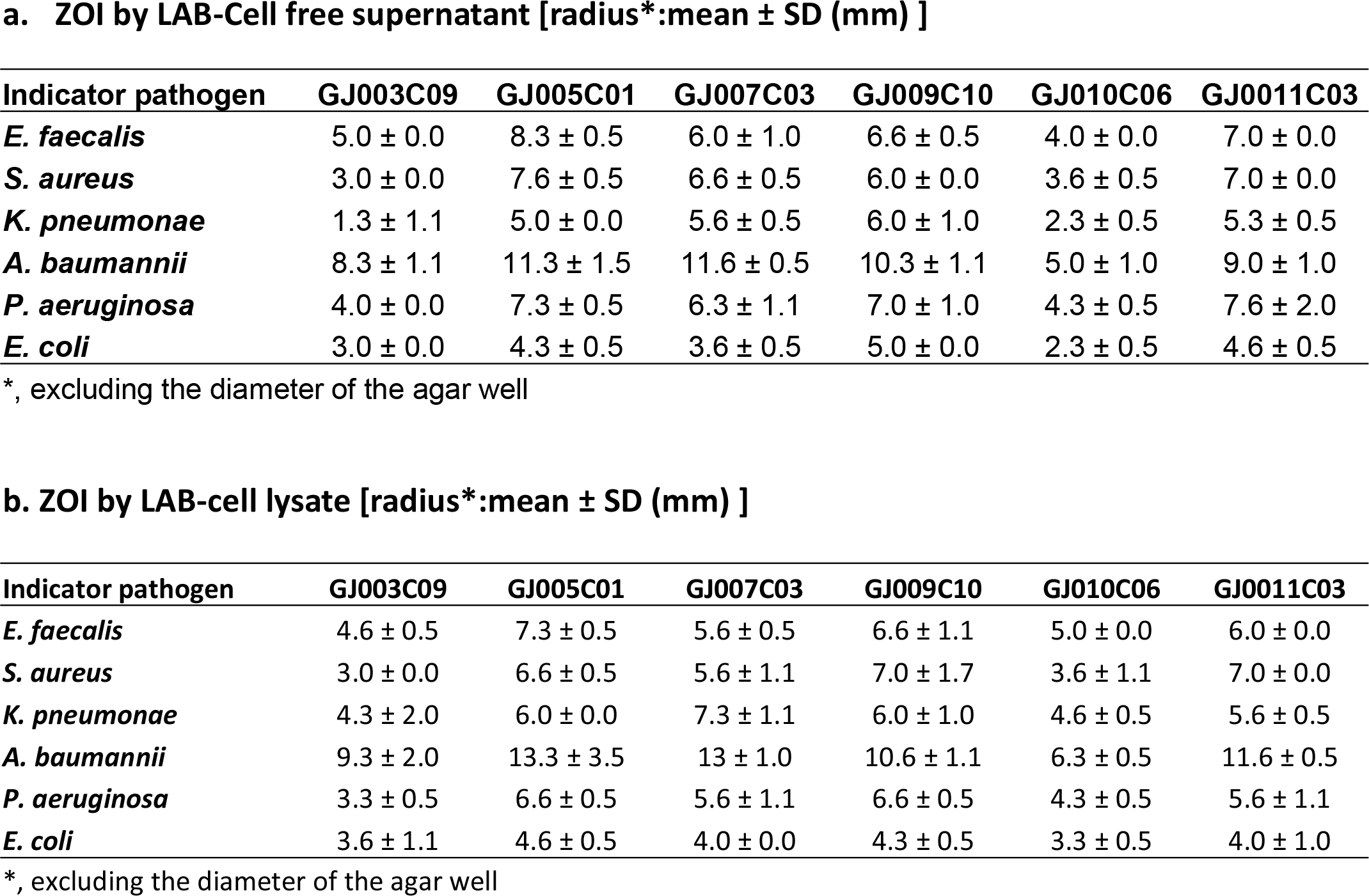
Zone of inhibition of *ESKAPE* pathogen growth by cell-free supernatant and cell-lysates of LAB isolates. The table depicts the Mean ± SD of the zone of inhibition radius from three consecutive experiments. The zone of inhibition was recorded with the zone of inhibition scale (PW297, Himedia) following 24 hours of incubation after the addition of sterile-filtered cell-free culture supernatants (**a**), and cell lysate (**b**) of LAB isolates to the agar-wells. All CSF and cell lysate rendered various levels of inhibition of ESKAPE pathogen growth of agar plates.

Cell-free supernatants of LAB isolates were found to exhibit anti-biofilm activity against ESKAPE pathogens (**Fig. 6**). All the LAB culture-free supernatants exhibited highly significant inhibition of biofilm formation in the cases of *E, faecalis*, *S. aureus*, *A. K. pneumoniae, A. baumannii,* and *P. aeruginosa,* although there is a certain degree of variation in their efficiency of inhibition (**Fig. 6 a-e**). In the case of *E. coli* biofilm, while the degree of inhibition is lower than that observed against the other five bacteria, the LAB- CFS of GJ007C03 and GJ011C03 afforded considerable inhibition, followed by moderate biofilm inhibition observed with CFS of GJ003C01, and GJ009C10 (**Fig. 6f**). Out of the 6 LAB isolates, the CSF of four isolates, specifically GJ009C10, GJ011C03, GJ005C01, and GJ007C03 significantly reduced the biofilm formation of *ESKAPE* pathogens.

**Fig. 6.**
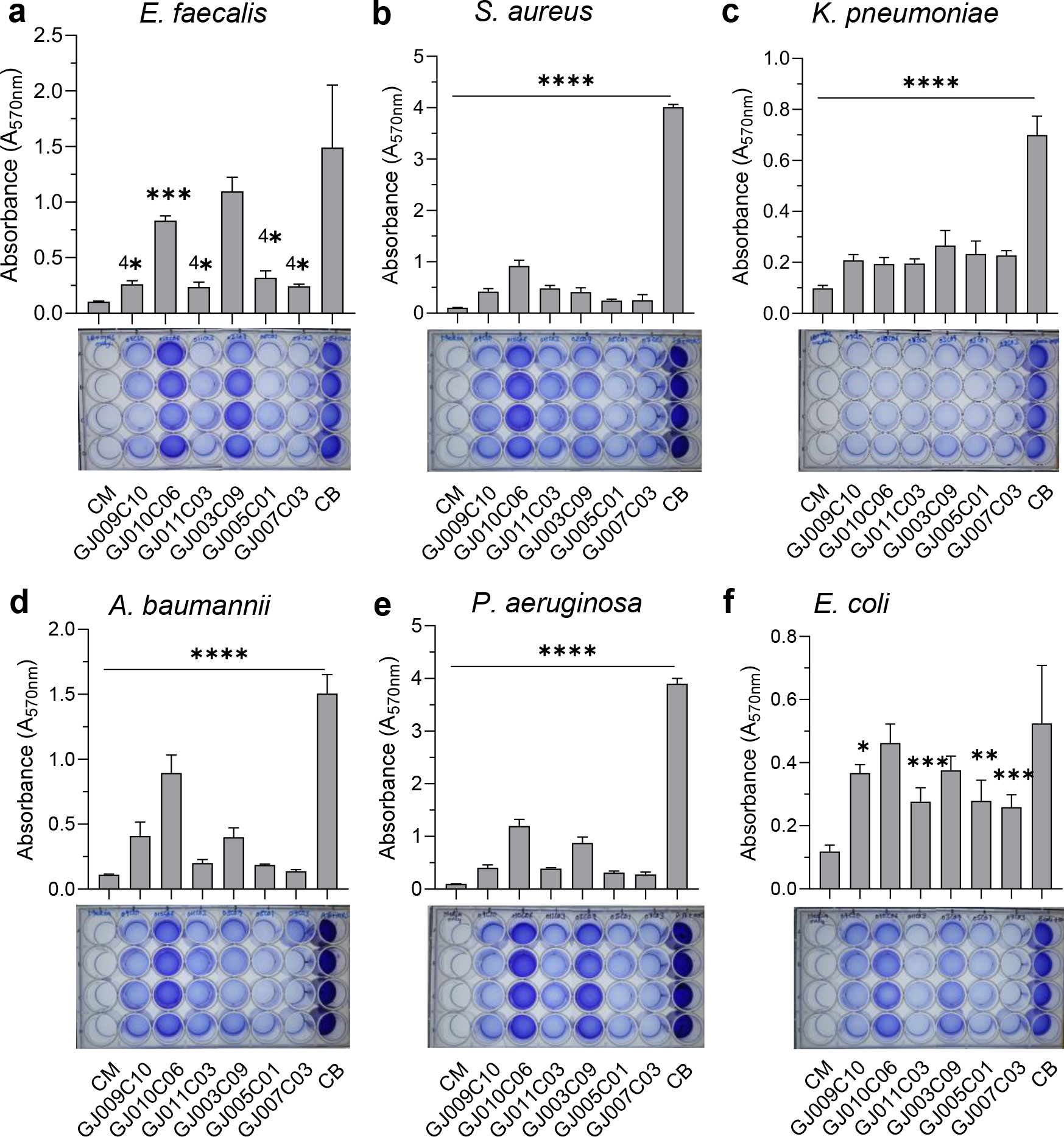
LAB isolates prevent biofilm formation by *ESKAPE* pathogens. The biofilm inhibition ability of LAB isolates was assessed using Cell-free culture filtrate (CSF). The assay was performed in 24-well polystyrene plates. ESKAPE pathogens were cultured either in the presence or absence of LAB-CSF and incubated in static conditions at 37 °C for 48 hours. Subsequently, quantitative measurement of biofilm was performed using the standard crystal violet-based assay. The bar diagrams depict the Mean ± SD of the absorbance (A_570nm_) values from triplicate wells. The experiment was performed twice. One-way ANOVA was performed to compare the mean differences of each condition compared to the base value without any CSF (control biofilm, CB). *, p<.05, **, p<.01, ***, p<.001, ****, p<.0001. All LAB-CSF exhibited various degrees of biofilm inhibition on ESKAPE bacteria.

## Discussion

The use of probiotics for animals has gained significant interest in recent years due to their numerous health benefits and market demands. While most commercial probiotics used in livestock are of human or dairy origin, probiotics from a similar host origin as the target host are preferred for their greater evolutionary adaptability with the host’s gastrointestinal environment and superior biological activity [63]. Probiotics isolated from animal intestines possess distinct characteristics, such as higher resistance to bile salts and low pH levels, as well as enhanced intestinal adhering abilities compared to dairy-based probiotics [64, 65]. Additionally, there is a growing need to develop species-specific probiotics capable of combating DR and MDR pathogens for improved health and performance of livestock animals [66, 67]. Previous studies have reported the isolation and characterization of probiotic bacterial species from goat milk and feces [68, 69]; however, there are currently no reports on LAB species specifically derived from the jejunum intestinal compartment. Thus, this study aimed to isolate and characterize Lactobacillus species from goat small intestine-jejunum, focusing on their morphological, biochemical, molecular, functional, and antagonistic properties against the proliferation and biofilm formation by the *ESKAPE* group of pathogens. The selection of probiotic bacteria was based on various criteria outlined by the Food and Agriculture Organization (FAO) and World Health Organization (WHO) [70]. These criteria include bacterial origin, species, strain characterization, functional aspects (gastric acid and bile tolerance, carbohydrate utilization, antimicrobial activity), antibiotic resistance, surface properties, and biosafety assessment [70]. Here out of 54 morphologically similar prospective LAB colonies grew on MRS agar from the goat-jejunum tissue homogenates, following multi-layered and multi-parametric identification via molecular, microbiological, and biochemical methods, only 6 isolates that were gram-positive, bacillus, positive for esculin hydrolysis, and negative for catalase, indole production, nitrate utilization, Voges Proskauer test, and Methyl Red tests, could ferment complex carbohydrates and showed > 99% sequence similarity with 16S rRNA gene, and 16S-23S ISR region for *Lactobacillus spp.* were shortlisted for further assessment of beneficial probiotic properties.

Probiotic candidates should tolerate low pH and bile salts to survive in the gastrointestinal tract [46]. Bacteria must survive the low pH of < 2.0 in the stomach in the case of simple stomach animals and < 2.5 in the abomasum in the case of ruminants and remain viable for at least ∼4 hours before reaching the intestines. Further, free bile acid that is synthesized in the liver conjugates with amino acids glycine or taurine, generating conjugated bile salts and releasing them in the duodenum [71]. These bile salts render antimicrobial activity by damaging bacterial cell walls and inducing DNA damage [72]. To cope with bile salts, bacteria in the gut must possess intrinsic resistance mechanisms [73]. Certain probiotic strains, including *Lactobacillus spp*, have specific proteins devoted to the efflux of bile salts and protons and modifying sugar metabolism, and preventing misfolding of proteins [73]. In this study, the survival rates of different LAB isolates varied under acidic and bile salt conditions. Some strains showed considerable survival at pH 2.0 and 3.0, while others demonstrated tolerance to different concentrations of bile salts, suggesting a strain-specific pattern. Notably, maximum survival or tolerance to acidity was observed in the case of GJ011C06 at lower pH, and GJ009C10, GJ010C06, and GJ011C03 tolerated higher bile salt environment relatively better than other three LAB isolates.

To increase the chances of survival and colonization in the gastrointestinal tract, probiotic bacteria must adhere to the intestinal epithelium [52]. Both auto-aggregation and surface hydrophobicity testing serve as pre-tests for evaluating the adhesion capacity of probiotic bacteria to epithelial cells. Various structures and components of bacteria, such as pili, fimbriae, adhesins, mucus-binding proteins, fibronectin-binding proteins, or surface layer proteins, lipoteichoic acid, and exopolysaccharides provides epithelial colonization enable epithelial colonization [49]. This attachment serves a protective role by competing with the intestinal pathobionts for host cell binding sites. All the LAB isolates in our study exhibited hydrophobicity values ranging from 60% to 80% by both the Xylene and Hexane methods. Similar hydrophobicity values were reported in several earlier studies for LAB isolates [74], and >40% hydrophobicity showing LABS were selected for feed supplement in the case of swine [63]. Further, the LAB isolates exhibited significant auto-aggregation ranging from 60-80%, indicating their potential for colonization and adhesion in the gastrointestinal tract.

Microbial communities as biofilms, when attached to a substratum or each other, are effective in controlling biofilm formation by other bacterial species [26]. *Lactobacillus spp.* known to form robust biofilms compared to other species of bacteria [54]. Probiotic biofilms can enhance colonization, prolong persistence in the intestinal mucosa, and regulate pathogenic biofilms [21]. In this study, the LAB isolates exhibited varied levels of biofilm formation under both aerobic and anaerobic conditions, however with significantly higher levels of biofilms in the anaerobic condition, and the top three biofilm-forming LAB isolates were GJ005C01, GJ009C10, and GJ011C03. In addition to the anti-pathobionts effect and increased colonization due to enhanced biofilm formation by probiotic LABs in the anaerobic condition in the gut, it may exert other beneficial effects. These include (i) enhanced nutrient absorption: biofilms provide a protective layer and retain nutrients that may be lost in the conventional gastrointestinal tract, allowing better nutrient absorption by the probiotic bacteria; (ii) defense against antibiotics: this enables the probiotic bacteria to survive and flourish even in the presence of antibiotics, thereby maintaining their beneficial effects on the gut during antibiotic therapy; and (iii) biofilm-mediated communication: biofilms are known to facilitate communication among bacteria, allowing for better coordination of metabolic and physiological functions and enabling the probiotic bacteria to function more efficiently for a better gut-homeostasis [75].

LABs can acquire as well as spread antibiotic resistance genes through horizontal gene transfer. Hence, antibiotic resistance testing is crucial when using new probiotic isolates as feed supplements due to the potential risk of transferring antibiotic resistance to pathogenic bacteria and reducing the efficacy of antibiotics. As per EU regulatory frameworks, any bacterial strain carrying an acquired gene conferring AMR or strains with the unknown genetic nature of a demonstrated resistance to antimicrobial agents should not be used as a feed additive due to the risk of horizontal spread [76]. In this study, all caprine gut-probiotic LAB isolates were found susceptible to all the antibiotics listed in the EFSA panel except vancomycin which may stems from the intrinsic resistance to vancomycin common in *Lactobacillus spp* [60, 77]. These findings on the selected LAB isolates, in addition to the non-hemolytic nature, ensure potential use as feed supplements to promote animal health and reduce antibiotic resistance.

All the six *ESKAPE* groups of pathogens commonly infect animals, with *Enterobacter* being the most common, followed by *K. pneumoniae, P. aeruginosa, S. aureus, E. faecium, and A. baumannii*. Major challenges of AMR in these pathogens are due to the multimodal antibiotic subversion mechanisms evolved by them, and biofilm formation is one of the leading causes. Therefore, agents that prevent biofilm formation and disperse the preformed biofilms are associated with therapeutic benefits to animals. Components of CFS of lactobacilli, such as, exopolysaccharides and biosurfactants, may inhibit the biofilm formation as previously reported against MDR pathogens [78, 79]. Anti-biofilm effect of CFS of *L. rhamnosus and L. plantarum* is well-known against food pathogens such as *P. aeruginosa* and *L. monocytogenes* [79]. Probiotic *Lactobacillus spp*. has also showed inhibition of biofilm formation as well as reduction in the gene expression involved in the quorum sensing pathway in *Streptococcus mutans* [80]. Anti-biofilm effect of cell-free supernatants of *L. pentosus* and *L. plantarum* were also reported against *B. cereus* and *P. aeruginosa* [81]. In this study, CFS of all six LAB isolates from goat small intestines inhibited biofilm formation of ESKAPE pathogens, with the variation in inhibition ability among the strains. Present findings are in agreement with a previous report for *Lactobacillus spp.* isolated from the GI tract of the other animal species [63].

Overall, the results of our study led to a preliminary depiction aimed at establishing a species-specific probiotic through the application of valid selection criteria. A total of six Lactobacillus isolates from the goat intestine were chosen for probiotic characterization. Specifically, the isolates examined in this study, namely GJ005C01, GJ007C03, GJ009C10, and GJ011C03, exhibited favorable characteristics such as robust survival in harsh gastric conditions (such as low pH and bile salts), biofilm formation, high hydrophobicity, and autoaggregation. None of the isolates displayed hemolytic activity, and all remained susceptible to nine antibiotics within the cut-off limits set by EFSA. Furthermore, all six Lactobacillus isolates demonstrated broad-spectrum antimicrobial activity and exhibited biofilm-inhibitory properties against ESKAPE pathogens. Among these isolates, GJ005C01, GJ007C03, GJ009C10, and

GJ011C03 showcased the most potent antibiofilm effects against ESKAPE pathogens and associated diseases. Therefore, they hold potential as livestock probiotics, serving as promising alternatives to antibiotics, and as next-generation probiotic candidates they could find utility in the feed and pharmaceutical industry.

## Materials and methods

### Ethics statement

All the experiments were reviewed and approved by the Institutional Biological Safety Committee (IBSC, Approval No. IBSC/2022/NIAB/BD/02) of the National Institute of Animal Biotechnology, Hyderabad, and all the procedure were performed by the relevant guidelines and regulations laid down by DBT-RCGM, Govt. of India. The requirement of the Institutional Animal Ethics Committee (IAEC) was waived as no animal experiments were involved, and only authorized slaughterhouse goat tissue samples were used for the isolation of probiotics.

### Bacterial strains and growth conditions

The *Lactobacillus plantarum*-MTCC strain that was used for proof of principle tests was procured from Microbial Type Culture Collection (MTCC), CSIR-Institute of Microbial Technology (IMTECH), Chandigarh, India. *LAB* isolates were grown in MRS agar or broth as applicable. Pathogenic *ESKAPE* group bacteria were obtained from: *E. coli*, and *S. aureus* (American type culture collection-ATCC, USA), *K. pneumonae*, *A. baumanii*, and *P. aeruginosa (MTCC, Chandigarh, India),* and *E. faecalis (National Center for Microbial Resources-NCMR, Pune, India)*. All bacteria were sub-cultured from glycerol stocks in Luria Bertani (LB) broth at 37°C overnight in a shaker incubator. Frozen glycerol stocks were prepared and stored at -80°C for future use.

### Isolation of Lactic Acid bacteria from goat intestinal tissue

Intestinal tissues (jejunum) were collected from goats (n = 11) immediately after slaughter in the Govt. authorized slaughterhouse in chilled phosphate buffer saline (PBS, pH 7.4), transported on ice to the laboratory, and processed on the same day for bacterial isolation in a BSL-2 facility. Goat intestinal tissue samples were thoroughly washed with 1XPBS to remove intestinal content and debris. Each intestinal tissue sample was dissected into small pieces and collected in screw-capped 2 ml homogenization tubes containing 1.0 mm glass beads. Tissue sections were homogenized two times for 30 seconds at 4000 rpm using a bead beater (BeadBug™). Subsequently, 100μl of homogenate was inoculated into 10 ml of sterile MRS broth. In parallel, intestinal content was collected separately in a 1.5 ml tube, and 100 μl of the soup was directly inoculated into 10 ml of MRS broth. All samples were incubated at 37°C in an orbital shaker incubator at 200 rpm for 24 hours. Subsequently, cultures were 10-fold serially diluted up to 1::10^6^, and 100ul samples from the three highest dilutions were plated onto MRS agar plates and incubated at 37°C aerobically for 48 hours. Single isolated colonies were picked and re-plated onto MRS agar for colony morphology evaluation and subsequent characterization.

### Biochemical characterization

Each colony was screened for colony and bacterial morphology, Gram staining, and catalase activity. Only Gram-positive and catalase-negative rods were further characterized. Various other tests were performed for the characterization of LAB as recommended in Bergey’s Manual of Determinative Bacteriology, including indole test, for the production of indole from tryptophan; nitrate test for the reduction of nitrate to nitrite; and Vogues Proskauer (VP) test to determine the presence of acetyl methyl carbinol after glucose fermentation [43]. The carbohydrate fermentation ability of all isolates was assessed by using the Himedia Hilacto identification kit (KB020, Himedia), which is a standardized, colorimetric identification system based on the principle of pH change and substrate utilization for the genus Lactobacillus. Kit contains one strip that includes 12 wells, one for esculin and another one for catalase, and 10 wells for 10 different carbohydrate sugars, i.e. Xylose, Cellobiose, Arabinose, Maltose, Galactose, Mannose, Mellibiose, Raffinose, Sucrose, and Trehalose. 50 μl of 0.1 OD bacterial inoculum was added to each well of a strip by surface inoculation method and incubated at 37°C for 24 - 48 hours. For carbohydrate fermentation and esculin test, the color change was observed in each well according to the manufacturer’s protocol. For the catalase test, a loopful of growth was scraped from the surface of the plate and dipped in a clean glass test tube containing 3% of freshly prepared H_2_O_2_, and effervescence was observed from the surface of the loop. No effervescence should be observed in case of a negative catalase test.

### Molecular identification of *LAB* isolates

For identification of the genus of the LAB isolates, colony-PCR was carried out on the genomic DNA extracted from the LAB isolates through the Triton-X-100 boiling lysis method using previously reported *Lactobacillus* genus-specific primers: Forward-R16-1, and Reverse-LbLMA1 as described previously (**Supplementary Table-1**) [38]. The optimized PCR cycle conditions included initial denaturation at 94°C for 1 minute, followed by 30 cycles consisting of denaturation at 98°C for 5 seconds, annealing at 55°C for 5 seconds, and extension at 72°C for 5 seconds, and 2 minutes of final extension. The amplified PCR products were analyzed using agarose gel electrophoresis with ethidium bromide staining and visualized under UV light.

For LAB species identification, a second round of PCR was performed targeting the 16S-rRNA V1-V3 region and 16S-23S Intergenic Spacer Region (ISR). Two separate primer pair sets were used to amplify ∼509 bp, and ∼565 bp, respectively using 16S(8-27)-F and V3(519-536)-R, and 16S(ISR)-F and 23S(ISR)-R primer pairs, respectively (**Supplementary Table-1**) [39–42]. The first PCR was conducted with sequencing primer set-1 under the following conditions: initial denaturation at 95°C for 10 minutes, followed by 40 cycles of denaturation at 95°C for 30 seconds, annealing at 57°C for 30 seconds, extension at 72°C for 60 seconds, and final extension of 7 minutes. For primer set-2, PCR conditions involved an initial denaturation at 95°C for 10 minutes, followed by 40 cycles of denaturation at 95°C for 30 seconds, annealing at 59°C for 30 seconds, extension at 72°C for 60 seconds, and a final extension of 7 minutes. The amplified PCR products were subjected to agarose gel electrophoresis and ethidium bromide staining. PCR products were subsequently excised from the agarose gel, and DNA was extracted using the Qiagen gel extraction kit (#28704) according to the manufacturer’s protocol. The excised amplicons were sequenced via Sanger sequencing using the forward primers of each PCR described above. The trimmed sequences were subjected to the NCBI-BLAST analysis with default parameters for identifying the closest bacterial species via % sequence cover and percentage identity.

Subsequently, phylogenetic analysis of the 12 LAB isolates was performed using either of the sequences of the 16S rRNA-V1-V3 region or 16S-23S ISR via hierarchical clustering using the Neighbour Joining method. Briefly, both sequences were aligned using ClustalW (default parameter). and the alignment results were then used to find out the best-fitted nucleotide-substitutional model. The Kimura 2- parameter model and Tamura 3-parameter model were used for the V3 region and the ISR region, respectively. Neighbor-joining method (bootstrap test -1000 replicates) was used to generate phylogenetic trees. Evolutionary analyses were conducted in MEGA11.

### Acid and bile tolerance test

The pH or acid tolerance of the samples was assessed under three different conditions: pH 2.0, 3.0, and 4.0, by adjusting the pH of MRS growth. In brief, 96-well microplates were added with 150 μl/well of MRS broth, adjusted to three different pH levels, and inoculated with 1% (V/V) bacterial culture from an overnight broth culture, which had been adjusted to 0.8 OD (A_600nm_). The microplates were then incubated at 37°C for 12 hours at 200 rpm in a microplate incubator. Absorbance was read at 600nm at two-hour intervals for a total of 12 hours.

To evaluate the resistance of LAB isolates in high bile salt conditions, the isolates were cultured in MRS broth containing three different bile salt concentrations: 0.5%, 1%, and 2% w/v. The same culture method was employed as described above for pH shock, with the exception that standard MRS media (pH 5.5) was used as the base media along with the appropriate bile salt concentration.

### Cell surface hydrophobicity assay

The cell surface hydrophobicity of Lactobacillus isolates was determined using the microbial adhesion to hydrocarbons (MATH) method as previously described [51]. This method evaluates hydrophobicity by measuring the affinity of microorganisms to an organic solvent such as hexane, xylene, or toluene. The LAB isolates were grown overnight in a shaker incubator at 37°C in MRS broth and then harvested by centrifugation at 8000 rpm at room temperature. The cells were washed twice using 1 x PBS and adjusted to an optical density of 0.8–1.0 at A_600nm_ (A_0_). Next, 1 ml of xylene or hexane was added to each suspension, and the mixture was vortexed vigorously for 2 min. After 1 hour of incubation at room temperature without shaking and phase separation, the aqueous phase was carefully removed, and its absorbance (A_t_) was measured. The hydrophobicity percentage was calculated using the following formula: Hydrophobicity (%) = (1- A_t_/A_0_) x 100.

### Evaluation of auto-aggregation

Auto-aggregation assays were performed as described previously [82]. Briefly, *Lactobacillus* isolates were grown overnight in MRS broth at 37°C and harvested by centrifugation at 8000 rpm at room temperature. The cells were then washed twice with 1 x PBS (pH= 7.4), and the resulting bacterials suspensions’ OD was adjusted to 0.8-1.0 at A_600nm_ (A_0_). They were subsequently incubated for 2 hours at 37°C without shaking. The upper phase was removed, and its OD was measured (A_t_). Finally, the auto-aggregation percentage was determined using the following formula: Auto−aggregation (%) = (OD A_0_ − OD At /A_0_) × 100.

### Biofilm formation assay

The biofilm formation ability of LAB isolates was assessed using a previously described protocol with minor modifications [83]. The test was carried out under both aerobic and anaerobic conditions. Briefly, *Lactobacillus* isolates were cultured overnight in MRS broth at 37 °C in a shaker incubator from glycerol stock to obtain primary culture. After this, all isolates were subcultured to achieve log phase growth (0.5-0.6 OD at A_600nm_). In each well of a 24-well polystyrene plate, 100 μl of secondary log phase culture was added to 2 ml of MRS medium and incubated at 37 °C for 72 hours. After incubation, non-adhered bacteria, and culture medium were eliminated, and the wells were washed twice with sterile distilled water. Plastic-adhered biofilms were fixed with 1 ml of methanol for 15 minutes, then methanol was removed, and the biofilms were air dried. Finally, the biofilms were stained by using 200 µl of 0.2% crystal violet in distilled water for 10 minutes, the excess stain was removed with sterile distilled water, and the stain was extracted from the adherent cells using 500 µl of 0.5 M glacial acetic acid. The absorbance was measured in a microplate reader at A_570nm_.

### Hemolytic assay

Overnight grown *LAB* cultures were streaked onto blood agar media containing 5% goat blood and incubated at 37°C for 24 h. *Listeria monocytogenes* and *S. aureus* were used as positive controls for α- and β- hemolysis, respectively. The clear and colored zones surrounding the colonies were examined. Clear zones were indicative of beta hemolysis, greenish zones were indicative of alpha hemolysis, and the absence of zones indicated no hemolysis or gamma hemolysis.

### Measurement of antibiotic susceptibility and the Minimum inhibitory concentration (MIC)

The antibiotic susceptibility of the *LAB i*solates was tested using the broth microdilution method. Nine antibiotics were selected including b-lactams like ampicillin (0.125-64 mg/L), macrolides like erythromycin (0.0039-2 mg/L), lincosamides like clindamycin (0.0156-8 mg/L), glycopeptides like vancomycin (0.125-64 mg/L), aminoglycosides like gentamicin (0.031-16 mg/L), kanamycin (0.125-64 mg/L), streptomycin (0.125-64 mg/L), tetracycline (0.125-64 mg/L) and chloramphenicol (0.125-64mg/L) as recommended by FEEDAP and EFSA panel for human and veterinary importance [60]. In brief, the isolated *LABs* were propagated overnight in MRS broth at 37°C to reach an OD (A_600nm_) of 1.0. Microliter test plates containing 2-fold serially diluted antibiotics in LAB susceptibility medium (LSM-broth) were inoculated at a final inoculum density of 0.001 (A_600nm_), equivalent to 10^5^ CFU/ml. After aerobic incubation at 37 °C for 24 and 48 hours, MIC values of each antibiotics were visually evaluated two times at 24 and 48 hours as the lowest antibiotic concentrations at which no growth was observed. The susceptibility status of strains was interpreted per the microbiological cut-off values defined by the EFSA Panel on bacterial Feed Additives and Products or Substances used in Animal Feed in relation to antimicrobial resistance [84].

### Assessment of antimicrobial activity

The antimicrobial activity of all *LAB* isolates was analyzed against ESKAPE pathogens using the agar well diffusion assay. Six different pathogenic bacteria were included in the study: such as *E. Coli, A. baumannii, P. aeruginosa, K. pneumonae,* 1. *S. aureus, and E. faecalis.* ESKAPE bacteria were grown overnight up to an OD of 1.0, and 10-fold serial dilutions were made in normal saline. 100µl of the bacterial inoculum was spread onto nutrient agar plates. Wells of 10 mm diameter were made on the agar plates using 1 ml sterile tip bottoms. For extraction of CFS of *LAB* isolates, they were grown overnight and centrifuged at 9,000 rpm at 4°C, and supernatants were collected and filter sterilized with a 0.2-micron filter. 100µl of CFS was added to the respective well on each agar plate. To get bacterial cell extract, overnight cultures of LAB were centrifuged at 9000 rpm at 4°C, and the supernatant was discarded. Cell pellets were washed with 1x PBS (phosphate buffer saline, pH 7.4), and cells were disrupted by homogenization using a bead-beater using 0.1mm beads in phosphate buffer; finally, following centrifugation cell lysate was filtered through a 0.2-micron filter and the sterile filtrate was used for the antimicrobial assay. After 24 hours of incubation at 37°C in aerobic conditions, the zone inhibition (mm) was recorded with the Zone of inhibition scale (PW297, Himedia).

### Biofilm inhibition assay

The antibiofilm property of CFS of *LAB* isolates on *ESKAPE* pathogens was assessed in 24-well plates. Briefly, 500μl of *ESKAPE* bacterial culture (OD: 0.3) in LB broth was added to each well along with 500μl of CFS (in MRS media). 1::1 mixture of L.B broth and MRS was used as the negative control. *ESKAPE* bacterial culture (0.3 OD) without CFS was used as a positive control. The microliter plates were incubated at 37°C for 48 hours to allow biofilm formation. Subsequently, biofilm was measured as described above in the ‘Biofilm formation assay’ section.

### Statistical Analysis

GraphPad Prism 9 was used for the preparation of the graphs and to perform the statistical analysis. For comparison of group means, One-Way ANOVA or t-test was performed, and differences were considered significant when p<0.05. All the results are shown as the mean ± SD unless otherwise described in the corresponding figure legends.

## Acknowledgments

Financial support from the Department of Biotechnology (DBT)-NIAB intramural grant is thankfully acknowledged. Support by the Council of Scientific & Industrial Research (CSIR), University Grant Commission (UGC), and DBT, Govt. of India for providing JRF/SRF to PS, RK, and JRF to SKB, respectively, thankfully acknowledged.

## Author Contributions

Conceived and designed the experiments: BD, PS. Performed the experiments: PS, RA, RK. Analyzed the data: PS, SKB, BD. Contributed reagents/ materials/analysis tools/ facility: BD. Wrote the paper: PS, BD. Provided overall supervision throughout the study: BD.

## Supporting information

Please see the supplementary information file.

## Data availability

All the nucleotide sequence data are available in the NCBI Bio-project No. PRJNA985412, PRJNA986841, and PRJNA986842. All other data that support the findings of this study are available from the corresponding author upon reasonable request.

## References

1. Boucher, H.W., et al., Bad bugs, no drugs: no ESKAPE! An update from the Infectious Diseases Society of America. Clin Infect Dis, 2009. 48(1): p. 1–12.

2. Shankar, P.R., Book review: Tackling drug-resistant infections globally. Archives of Pharmacy Practice, 2016. 7(3): p. 110–111.

3. Mulani, M.S., et al., Emerging Strategies to Combat ESKAPE Pathogens in the Era of Antimicrobial Resistance: A Review. Front Microbiol, 2019. 10: p. 539.

4. Lewis, K., Persister cells and the riddle of biofilm survival. Biochemistry (Moscow), 2005. 70(2): p. 267–274.

5. Andersson, D.I., et al., Antibiotic resistance: turning evolutionary principles into clinical reality. FEMS Microbiology Reviews, 2020. 44(2): p. 171–188.

6. da Rosa, T.F., et al., Alternatives for the treatment of infections caused by ESKAPE pathogens. J Clin Pharm Ther, 2020. 45(4): p. 863–873.

7. Hill, C., et al., Expert consensus document: The International Scientific Association for Probiotics and Prebiotics consensus statement on the scope and appropriate use of the term probiotic. Nature reviews Gastroenterology & hepatology, 2014.

8. Mattia, A. and R. Merker, Regulation of probiotic substances as ingredients in foods: premarket approval or “generally recognized as safe” notification. Clinical infectious diseases, 2008. 46(Supplement_2): p. S115–S118.

9. Silva, D.R., et al., Probiotics as an alternative antimicrobial therapy: Current reality and future directions. Journal of Functional Foods, 2020. 73: p. 104080.

10. Bermudez-Brito, M., et al., Probiotic mechanisms of action. Ann Nutr Metab, 2012. 61(2): p. 160–74.

11. Stanbro, J., et al., Topical delivery of lactobacillus culture supernatant increases survival and wound resolution in traumatic Acinetobacter baumannii infections. Probiotics and antimicrobial proteins, 2020. 12: p. 809–818.

12. Jones, M., et al., Novel nitric oxide producing probiotic wound healing patch: preparation and in vivo analysis in a New Zealand white rabbit model of ischaemic and infected wounds. International wound journal, 2012. 9(3): p. 330–343.

13. Fijan, S., et al., Efficacy of Using Probiotics with Antagonistic Activity against Pathogens of Wound Infections: An Integrative Review of Literature. Biomed Res Int, 2019. 2019: p. 7585486.

14. Yilmaz, O. and S. Turkyilmaz, Investigation of the potential probiotic effects of lactic acid bacteria and cell-free supernatants against important pathogens leading to wound infections. 2022.

15. Velraeds, M.M., et al., Inhibition of initial adhesion of uropathogenic Enterococcus faecalis by biosurfactants from Lactobacillus isolates Appl Environ Microbiol, 1996. 62(6): p. 1958–63.

16. Rastogi, S., V. Mittal, and A. Singh, Selection of Potential Probiotic Bacteria from Exclusively Breastfed Infant Faeces with Antagonistic Activity Against Multidrug-Resistant ESKAPE Pathogens. Probiotics Antimicrob Proteins, 2021. 13(3): p. 739–750.

17. Fitzgerald, J.R., Livestock-associated Staphylococcus aureus: origin, evolution and public health threat. Trends Microbiol, 2012. 20(4): p. 192–8.

18. Kang, M.S., et al., Antimicrobial activity of Lactobacillus salivarius and Lactobacillus fermentum against Staphylococcus aureus. Pathog Dis, 2017. 75(2).

19. Munoz, M.A., et al., Fecal shedding of Klebsiella pneumoniae by dairy cows. J Dairy Sci, 2006. 89(9): p. 3425–30.

20. Darniati, D., et al., First evidence of Klebsiella pneumoniae infection in Aceh cattle: Pathomorphology and antigenic distribution in the lungs. Vet World, 2021. 14(4): p. 1007–1013.

21. Lagrafeuille, R., et al., Opposing effect of Lactobacillus on in vitro Klebsiella pneumoniae in biofilm and in an in vivo intestinal colonisation model. Benef Microbes, 2018. 9(1): p. 87–100.

22. Mogna, L., et al., In Vitro Inhibition of Klebsiella pneumoniae by Lactobacillus delbrueckii Subsp. delbrueckii LDD01 (DSM 22106): An Innovative Strategy to Possibly Counteract Such Infections in Humans? J Clin Gastroenterol, 2016. p. S136–s139.

23. Haenni, M., et al., Population structure and antimicrobial susceptibility of Pseudomonas aeruginosa from animal infections in France. BMC Vet Res, 2015. 11: p. 9.

24. Leitner, G. and O. Krifucks, Pseudomonas aeruginosa mastitis outbreaks in sheep and goat flocks: antibody production and vaccination in a mouse model. Vet Immunol Immunopathol, 2007. 119(3-4): p. 198–203.

25. Shokri, D., et al., The Inhibition Effect of Lactobacilli Against Growth and Biofilm Formation of Pseudomonas aeruginosa. Probiotics Antimicrob Proteins, 2018. 10(1): p. 34–42.

26. A. Elbadri, S., et al., The effect of Lactobacillus acidophilus as a probiotic against Pseudomonas aeruginosa growth and biofilm formation. Novel Research in Microbiology Journal, 2019. 3(4): p. 428–439.

27. Alexandre, Y., et al., Screening of Lactobacillus spp. for the prevention of Pseudomonas aeruginosa pulmonary infections. BMC Microbiology, 2014. 14(1): p. 107.

28. Health, E.P.o.A., et al., Assessment of listing and categorisation of animal diseases within the framework of the Animal Health Law (Regulation (EU) No 2016/429): antimicrobial-resistant Enterococcus faecalis in poultry. EFSA Journal, 2022. 20(2): p. e07127.

29. Hammerum, A.M., Enterococci of animal origin and their significance for public health. Clin Microbiol Infect, 2012. 18(7): p. 619–25.

30. Safadi, S., et al., The Products of Probiotic Bacteria Effectively Treat Persistent Enterococcus faecalis Biofilms. Pharmaceutics, 2022. 14(4): p. 751.

31. Kim, A.R., et al., Lactobacillus plantarum lipoteichoic acid disrupts mature Enterococcus faecalis biofilm. J Microbiol, 2020. 58(4): p. 314–319.

32. Frank, J.F. and E.H. Marth, Inhibition of Enteropathogenic Escherichia coli by Homofermentative Lactic Acid Bacteria in Skimmilk. J Food Prot, 1977. 40(11): p. 754–759.

33. Byakika, S., et al., Antimicrobial Activity of Lactic Acid Bacteria Starters against Acid Tolerant, Antibiotic Resistant, and Potentially Virulent E. coli Isolated from a Fermented Sorghum-Millet Beverage. Int J Microbiol, 2019. 2019: p. 2013539.

34. Campana, R., S. van Hemert, and W. Baffone, Strain-specific probiotic properties of lactic acid bacteria and their interference with human intestinal pathogens invasion. Gut pathogens, 2017. 9(1): p. 1–12.

35. Ramos, C.L., et al., Strain-specific probiotics properties of Lactobacillus fermentum, Lactobacillus plantarum and Lactobacillus brevis isolates from Brazilian food products. Food microbiology, 2013. 36(1): p. 22–29.

36. Dave, R. and N. Shah, Evaluation of media for selective enumeration of Streptococcus thermophilus, Lactobacillus delbrueckii ssp. bulgaricus, Lactobacillus acidophilus, and bifidobacteria. Journal of dairy science, 1996. 79(9): p. 1529–1536.

37. De Man, J., d. Rogosa, and M.E. Sharpe, A medium for the cultivation of lactobacilli. Journal of Applied Microbiology, 1960. 23(1): p. 130–135.

38. Dubernet, S., N. Desmasures, and M. Guéguen, A PCR-based method for identification of lactobacilli at the genus level. FEMS Microbiol Lett, 2002. 214(2): p. 271–5.

39. Berthier, R., et al., Adhesion of mature polyploid megakaryocytes to fibronectin is mediated by beta 1 integrins and leads to cell damage. Exp Cell Res, 1998. 242(1): p. 315–27.

40. Gurtler, V. and V.A. Stanisich, New approaches to typing and identification of bacteria using the 16S-23S rDNA spacer region. Microbiology (Reading), 1996. 142 (Pt 1): p. 3–16.

41. Weisburg, W.G., et al., 16S ribosomal DNA amplification for phylogenetic study. J Bacteriol, 1991. 173(2): p. 697–703.

42. Turner, S., et al., Investigating deep phylogenetic relationships among cyanobacteria and plastids by small subunit rRNA sequence analysis. J Eukaryot Microbiol, 1999. 46(4): p. 327–38.

43. Bergey, D.H., Bergey’s manual of determinative bacteriology. 1994: Lippincott Williams & Wilkins.

44. Matthews, C., et al., The rumen microbiome: a crucial consideration when optimising milk and meat production and nitrogen utilisation efficiency. Gut Microbes, 2019. 10(2): p. 115–132.

45. Chuard, C. and L.B. Reller, Bile-esculin test for presumptive identification of enterococci and streptococci: effects of bile concentration, inoculation technique, and incubation time. J Clin Microbiol, 1998. 36(4): p. 1135–6.

46. Chou, L.-S. and B. Weimer, Isolation and characterization of acid-and bile-tolerant isolates from strains of Lactobacillus acidophilus. Journal of Dairy Science, 1999. 82(1): p. 23–31.

47. Tuomola, E., et al., Quality assurance criteria for probiotic bacteria. The American journal of clinical nutrition, 2001. 73(2): p. 393s–398s.

48. Ohland, C.L. and W.K. MacNaughton, Probiotic bacteria and intestinal epithelial barrier function. American journal of physiology-gastrointestinal and liver physiology, 2010. 298(6): p. G807–G819.

49. Krausova, G., I. Hyrslova, and I. Hynstova, In vitro evaluation of adhesion capacity, hydrophobicity, and auto-aggregation of newly isolated potential probiotic strains. Fermentation, 2019. 5(4): p. 100.

50. Magnusson, K.-E., et al., Non-specific and specific recognition mechanisms of bacterial and mammalian cell membranes. Journal Of Dispersion Science Andtechnology, 1985. 6(1): p. 69–89.

51. Vinderola, C.G., M. Medici, and G. Perdigón, Relationship between interaction sites in the gut, hydrophobicity, mucosal immunomodulating capacities and cell wall protein profiles in indigenous and exogenous bacteria. J Appl Microbiol, 2004. 96(2): p. 230–43.

52. Tuo, Y., et al., Aggregation and adhesion properties of 22 Lactobacillus strains. Journal of dairy science, 2013. 96(7): p. 4252–4257.

53. Salas-Jara, M.J., et al., Biofilm forming Lactobacillus: new challenges for the development of probiotics. Microorganisms, 2016. 4(3): p. 35.

54. Kubota, H., et al., Biofilm Formation by Lactic Acid Bacteria and Resistance to Environmental Stress. Journal of Bioscience and Bioengineering, 2008. 106(4): p. 381–386.

55. Naidu, K.S.B., J.K. Adam, and P. Govender, The use of probiotics and safety concerns: A review. 2012.

56. Tanaka, Y., et al., In Vitro Probiotic Characterization and Safety Assessment of Lactic Acid Bacteria Isolated from Raw Milk of Japanese-Saanen Goat (Capra hircus). Animals, 2022. 13(1): p. 7.

57. Mogrovejo, D.C., et al., Prevalence of Antimicrobial Resistance and Hemolytic Phenotypes in Culturable Arctic Bacteria. Front Microbiol, 2020. 11: p. 570.

58. Authority, E.F.S., Opinion of the Scientific Panel on additives and products or substances used in animal feed (FEEDAP) on the updating of the criteria used in the assessment of bacteria for resistance to antibiotics of human or veterinary importance. EFSA Journal, 2005. 3(6): p. 223.

59. Brashears, M.M., A. Amezquita, and D. Jaroni, Lactic acid bacteria and their uses in animal feeding to improve food safety. Advances in food and nutrition research, 2005. 50: p. 1–31.

60. Additives, E.P.o. and P.o.S.u.i.A. Feed, Guidance on the assessment of bacterial susceptibility to antimicrobials of human and veterinary importance. EFSA Journal, 2012. 10(6): p. 2740.

61. Manyi-Loh, C., et al., Antibiotic use in agriculture and its consequential resistance in environmental sources: potential public health implications. Molecules, 2018. 23(4): p. 795.

62. Plaza-Diaz, J., et al., Mechanisms of Action of Probiotics. Adv Nutr, 2019. 10(suppl_1): p. S49–S66.

63. Dowarah, R., et al., Selection and characterization of probiotic lactic acid bacteria and its impact on growth, nutrient digestibility, health and antioxidant status in weaned piglets. PLoS One, 2018. 13(3): p. e0192978.

64. Sornplang, P. and S. Piyadeatsoontorn, Probiotic isolates from unconventional sources: a review. J Anim Sci Technol, 2016. 58: p. 26.

65. Bazireh, H., et al., Isolation of novel probiotic Lactobacillus and Enterococcus strains from human salivary and fecal sources. Frontiers in microbiology, 2020. 11: p. 597946.

66. Blajman, J., et al., In vitro and in vivo screening of native lactic acid bacteria toward their selection as a probiotic in broiler chickens. Res Vet Sci, 2015. 101: p. 50–6.

67. Ripamonti, B., et al., Screening of species-specific lactic acid bacteria for veal calves multi-strain probiotic adjuncts. Anaerobe, 2011. 17(3): p. 97–105.

68. Setyawardani, T., et al., Identification and characterization of probiotic lactic acid bacteria isolated from indigenous goat milk. Animal Production, 2011. 13(1).

69. Andrada, E., et al., Ferulic Acid Esterase Producing Lactobacillus johnsonii from Goat Feces as Corn Silage Inoculants. Microorganisms, 2022. 10(9): p. 1732.

70. Bajagai, Y.S., et al., Probiotics in animal nutrition: production, impact and regulation. 2016: FAO.

71. Casadesús Pursals, J. and V.C. Urdaneta Páez, interactions between Bacteria and Bile Salts in the Gastrointestinal and Hepatobiliary Tracts. Frontiers in Medicine, 4 (163), 1–13., 2017.

72. Begley, M., C.G. Gahan, and C. Hill, The interaction between bacteria and bile. FEMS microbiology reviews, 2005. 29(4): p. 625–651.

73. Ruiz, L., A. Margolles, and B. Sánchez, Bile resistance mechanisms in Lactobacillus and Bifidobacterium. Frontiers in microbiology, 2013. 4: p. 396.

74. Gomaa, E.Z., Antimicrobial and anti-adhesive properties of biosurfactant produced by lactobacilli isolates, biofilm formation and aggregation ability. The journal of general and applied microbiology, 2013. 59(6): p. 425–436.

75. Barzegari, A., et al., The battle of probiotics and their derivatives against biofilms. Infection and drug resistance, 2020: p. 659–672.

76. Stefańska, I., et al., Antimicrobial susceptibility of lactic acid bacteria strains of potential use as feed additives-the basic safety and usefulness criterion. Frontiers in Veterinary Science, 2021. 8: p. 687071.

77. Tynkkynen, S., K.V. Singh, and P. Varmanen, Vancomycin resistance factor of Lactobacillus rhamnosus GG in relation to enterococcal vancomycin resistance (van) genes. International journal of food microbiology, 1998. 41(3): p. 195–204.

78. Kaur, S., et al., Anti-biofilm properties of the fecal probiotic lactobacilli against Vibrio spp. Frontiers in cellular and infection microbiology, 2018. 8: p. 120.

79. Rezaei, Z., S. Khanzadi, and A. Salari, Biofilm formation and antagonistic activity of Lacticaseibacillus rhamnosus (PTCC1712) and Lactiplantibacillus plantarum (PTCC1745). AMB Express, 2021. 11(1): p. 1–7.

80. Wasfi, R., et al., Probiotic Lactobacillus sp. inhibit growth, biofilm formation and gene expression of caries-inducing Streptococcus mutans. Journal of cellular and molecular medicine, 2018. 22(3): p. 1972–1983.

81. Khiralla, G.M., et al., Antibiofilm effect of Lactobacillus pentosus and Lactobacillus plantarum cell-free supernatants against some bacterial pathogens. Journal of Biotech Research, 2015. 6: p. 86.

82. Zuo, F., et al., Characterization and in vitro properties of potential probiotic Bifidobacterium strains isolated from breast-fed infant feces. Annals of Microbiology, 2015. 66.

83. Aoudia, N., et al., Biofilms of Lactobacillus plantarum and Lactobacillus fermentum: effect on stress responses, antagonistic effects on pathogen growth and immunomodulatory properties. Food Microbiology, 2016. 53: p. 51–59.

84. Hazards, E.P.o.B., et al., Update of the list of QPS-recommended biological agents intentionally added to food or feed as notified to EFSA 14: suitability of taxonomic units notified to EFSA until March 2021. EFSA Journal, 2021. 19(7): p. e06689.

