## Supplementary information for "Restriction of the growth and biofilm formation of *ESKAPE* pathogens by caprine gut-derived probiotic bacteria"

[3]current address: The University of Alabama in Huntsville, Chemical & Material Engineering, 301 Sparkman Drive, Huntsville, Alabama, USA 35899

***Corresponding address:**

Bappaditya Dey
Scientist-E, National Institute of Animal Biotechnology (NIAB)
Survey No. 37, Extended Q City Road, Gowlidoddi, Gachibowli
Hyderabad, Telangana, India 500032

Ph: +917042077704

**Supplementary Table-1: PCR primers used for identification of *Lactobacillus* isolates**

| Name | Nucleotide Sequence | Target | Reference |
|---|---|---|---|
| ***Lactobacillus* genus specific PCR primer** | | | |
| **R16-1** | CTTGTACACACCGCCCGTCA | 16s-rRNA gene | 38 |
| **LbLMA1-rev** | CTCAAAACTAAACAAAGTTTC | | |
| ***Lactobacillus sp.* PCR and sequencing primer** | | | |
| **16S(ISR)-F** | GCTGGATCACCTCCTTTC | 16S-23S ISR | 39-42 |
| **23S(ISR)-R** | CCTTTCCCTCACGGTACTG | | |
| **16S(8-27)-F** | AGAGTTTGATCCTGGCTCAG | 16-rRNA-V1-V3 region | |
| **V3(519-536)-R** | GTATTACCGCGGCTGCTG | | |

**Supplementary Fig. 1**

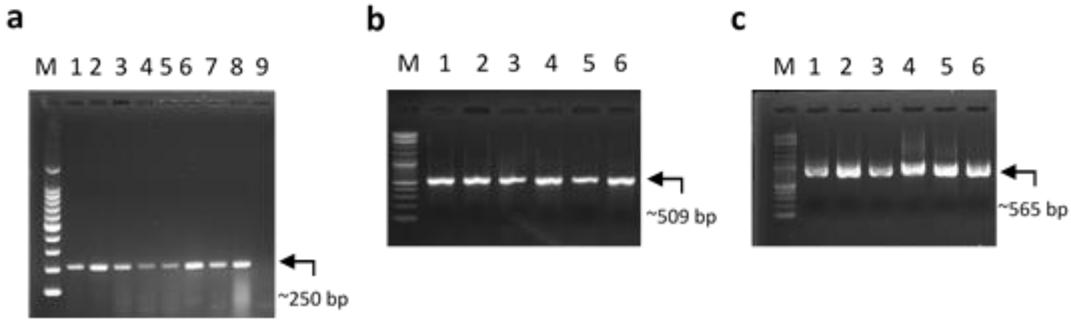


**Supplementary Fig. 1. PCR and agarose gel electrophoresis analysis for molecular identification of LAB isolates.**

Representative agarose gel electrophoresis analysis images of (**a**) *Lactobacillus* genus-specific PCR analysis using 16S-rRNA gene-specific primers, (**b**) 16S rRNA gene V1-V3 region, and (**c**) 16S-23S ISR of genomic DNA from LAB isolates. Following is the lane description.

a. 1. *L. plantarum* (+ve control), 2. *GJ003C09,* 3. GJ005C01, 4. GJ007C03, 5. GJ009C10, 6. GJ010C06, 7. GJ011C03, 8. GJ010C02, 9. PCR –ve control.

b. 1. *GJ003C09,* 2. GJ005C01, 3. GJ007C03, 4. GJ009C10, 5. GJ010C06, and 6. GJ011C03

c. 1. *GJ003C09,* 2. GJ005C01, 3. GJ007C03, 4. GJ009C10, 5. GJ010C06, and 6. GJ011C03

**Supplementary Fig. 2**

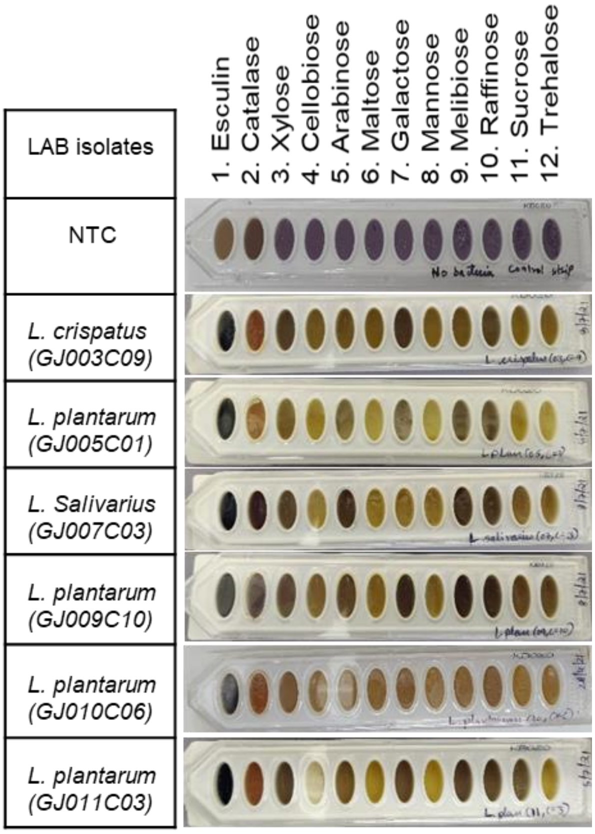


**Supplementary Fig. 2. Carbohydrate fermentation ability of LAB isolates.** The carbohydrate fermentation ability of LAB isolates was assessed by using the Hilacto identification kit (KB020, Himedia). One assay strip has 12 wells including 10 different carbohydrate sugars, one for esculin, and the other one for catalase. The carbohydrate fermentation was measured via colorimetric identification based on the principle of pH change and substrate utilization for the genus *Lactobacillus*. While dark coloration confirms esculin hydrolysis (**Lane -1**), light coloration indicates sugar fermentation (**Lane 3-12**). For the catalase test visual observation of effervescence was considered a positive test, and no effervescence for the negative catalase test (**Lane- 2**). All 6 LAB isolates were found positive for esculin, negative for catalase, and with varied abilities to ferment different types of sugar. NTC: Negative control.

**Supplementary Fig. 3**

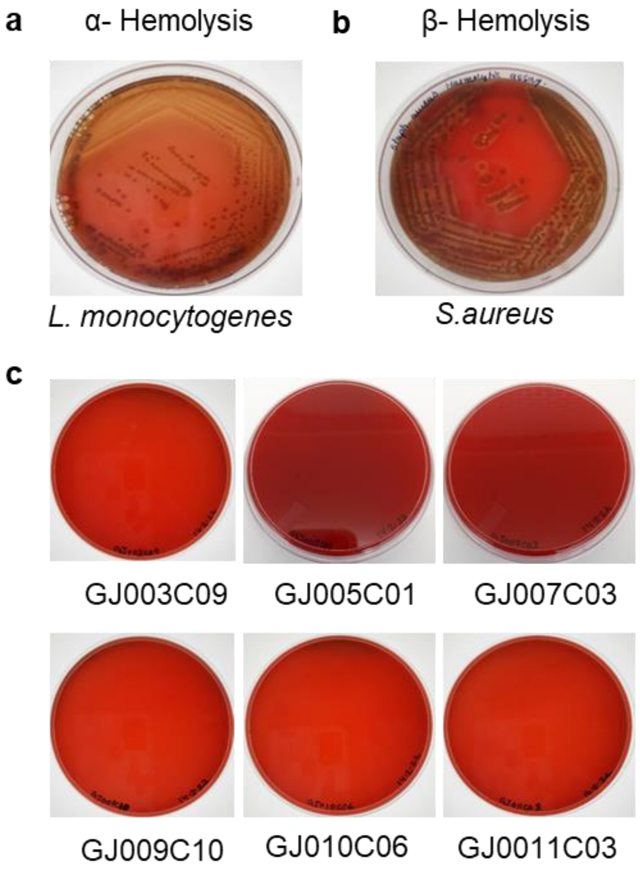


**Supplementary Fig. 3. LAB isolates are non-hemolytic**: Overnight-grown LAB cultures were streaked onto blood agar media containing 5% goat blood and incubated at 37˚C for 24 hours. Two positive control bacteria, Listeria monocytogenes, and Staphylococcus aureus, were used for α- and β- hemolysis respectively. The clear and colored zones surrounding the colonies were examined. Clear zones were indicative of beta hemolysis, greenish zones were indicative of alpha hemolysis and the absence of zones indicated no hemolysis or gamma hemolysis. All 6 LAB isolates did not show any hemolysis on the blood-agar plates.
